# *Trans*-interaction of risk loci 6p24.1 and 10q11.21 is associated with endothelial damage in coronary artery disease

**DOI:** 10.1101/2022.07.12.499721

**Authors:** Kai Yi Tay, Kan Xing Wu, Florence Wen Jing Chioh, Matias Ilmari Autio, Nicole Min Qian Pek, Balakrishnan Chakrapani Narmada, Sock-Hwee Tan, Adrian Fatt-Hoe Low, Michelle Mulan Lian, Elaine Guo Yan Chew, Hwee Hui Lau, Shih Ling Kao, Adrian Kee Keong Teo, Jia Nee Foo, Roger Sik Yin Foo, Chew Kiat Heng, Mark Yan Yee Chan, Christine Cheung

## Abstract

**Background and Aims:** Single nucleotide polymorphism rs6903956 has been identified as one of the genetic risk factors for coronary artery disease (CAD). However, rs6903956 lies in a non-coding locus on chromosome 6p24.1. We aim to interrogate the molecular basis of 6p24.1 containing rs6903956 risk alleles in endothelial disease biology.

**Methods and Results:** We generated induced pluripotent stem cells (iPSCs) from CAD patients (AA risk genotype at rs6903956) and normal controls (GG non-risk genotype at rs6903956). CRIPSR-Cas9-based deletions (Δ63-89bp) on 6p24.1, including both rs6903956 and a short tandem repeat variant rs140361069 in linkage disequilibrium, were performed to generate isogenic iPSC-derived endothelial cells. Edited CAD endothelial cells, with removal of ‘A’ risk alleles, exhibited a global transcriptional downregulation of pathways relating to abnormal vascular physiology and activated endothelial processes. A CXC chemokine ligand on chromosome 10q11.21, *CXCL12*, was uncovered as a potential effector gene in CAD endothelial cells. Underlying this effect was the preferential inter-chromosomal interaction of 6p24.1 risk locus to a weak promoter of *CXCL12*, confirmed by chromatin conformation capture assays on our iPSC-derived endothelial cells. Functionally, risk genotypes AA/ AG at rs6903956 were associated significantly with elevated levels of circulating damaged endothelial cells in CAD patients. Circulating endothelial cells isolated from patients with risk genotypes AA/ AG were also found to have 10 folds higher CXCL12 transcript copies/ cell than those with non-risk genotype GG.

**Conclusion:** Our study reveals the trans-acting impact of 6p24.1 with another CAD locus on 10q11.21 and is associated with intensified endothelial injury.

## Introduction

Cardiovascular disease is the leading cause of mortality globally, with coronary artery disease (CAD) accounting for a majority of cardiovascular deaths [1]. With the heritability of CAD estimated at up to 60% [2], genetic factors hold an important contribution to the risk of CAD along with other major etiologic determinants such as lifestyle and environmental factors. Inter-ethnic differences in the prevalence of CAD and cardiovascular risk factors have been observed [3, 4].

Genome-wide association studies (GWAS) have identified several common variants to be risk factors for CAD, but few have been functionally validated [5–7]. A single nucleotide polymorphism (SNP) rs6903956 on chromosome 6p24.1 was first reported as a susceptibility locus for CAD (adjusted odds ratio = 1.65, 95% CI 1.44-1.90, *p* = 2.55 × 10^-13^, Supplemental Table S1) in a GWAS on large cohort Chinese population [8]. The minor allele at rs6903956 (A, 9.07% frequency) leads to increased risk of coronary atherosclerosis (Supplemental Table S1) [9]. This finding has been subsequently corroborated by several independent studies on Asian populations, including Singaporean and Japanese individuals [9–12]. In our Singaporean population, we leveraged on the ‘Singapore Coronary Artery Disease Genetics Study’ where CAD and normal control subjects from the National University Hospital Angiography Centre had been genotyped for a CAD GWAS [13]. The rs6903956 variant was one of the constituent SNPs being interrogated in the genotyping array. We confirm that heterozygous AG at rs6903956 is significantly associated with CAD in Chinese (adjusted odds ratio = 2.26, 95% CI 1.09–4.6, *p* = 0.028), and not in the other ethnic groups when analyzed independently [11].

SNP rs6903956 lies within the first intron of androgen-dependent tissue factor pathway inhibitor regulating protein (*ADTRP*). Its minor risk allele A is associated with decreased *ADTRP* mRNA expression in leukocytes [8]. Plasma levels of ADTRP are significantly reduced in CAD patients compared to control subjects [14]. As risk variants identified by GWAS located in intronic regions of the genome are generally believed to confer risk of disease via disruption of gene regulatory elements [15], a recent study found that binding of GATA2 with a 519bp region containing rs6903956 non-risk allele G enhances *ADTRP* expression level in HeLa cells, suggesting a putative enhancer role of this region [16]. Genetic effects are often restricted to trait relevant cell types, making it important to have precise disease-specific tissue and cell types for expression quantitative trait loci (eQTL) mapping [17, 18]. However, there remains a lack of eQTL analysis of rs6903956 on relevant tissues from CAD patients.

CAD is largely caused by atherosclerosis, which is a build-up of plaque inside the artery walls. Multiple cell types are involved in CAD pathogenesis, including endothelial cells, vascular smooth muscle cells, macrophages, and cells of the adaptive immune system [19]. Functional causality of genetic risk variants could be better resolved with a focus on cell-type specific effects. For instance, the well-studied CAD risk locus 9p21.3 mediates its effect on *ANRIL* expression, and induced proinflammatory responses in endothelial cells, vascular smooth muscle cells and mononuclear cells [20–23]. A common noncoding SNP rs17114036 at 1p32.2 is located in an endothelial-specific enhancer and regulates endothelial mechanotransduction mechanisms [24]. Also, within our 6p24 locus of interest, another SNP rs9349379 in the third intron of gene encoding phosphatase and actin regulatory protein 1 (*PHACTR1*) demonstrates *cis*-regulation of endothelial expression of endothelin-1 (*EDN1*), a gene located 600kb upstream of *PHACTR1* [25]. It has been reported that ADTRP confers anti-coagulant protection in endothelial cells through regulation of tissue factor pathway inhibitor [26]. Here, we prioritized endothelial disease biology in interrogating the functional basis of 6p24 locus that contains rs6903956.

Given the challenges of obtaining patient coronary arteries and the tendency of primary endothelial cells to senesce *in vitro*, we leveraged induced pluripotent stem cells (iPSC) technology to generate human iPSC-derived endothelial cells. In analyzing nearby variants of rs6903956, a short tandem repeat variant rs140361069 was found to be the nearest variant in linkage disequilibrium (LD) with rs6903956 in East Asian populations. We postulated that the ethnicity-dependent effect of rs6903956 might partially be due to the lack of rs140361069 in LD with rs6903956 in Europeans and most of the other populations. Hence, we employed CRISPR-Cas9 genome editing to delete small regions (63 – 89bp) covering both rs6903956 and rs140361069 at 6p24.1 in CAD patient iPSCs and normal subject iPSCs. Using protocols that we had previously established [27, 28], isogenic iPSC lines were differentiated using chemically-defined factors to obtain consistent cultures of edited and unedited iPSC-derived endothelial cells. To decipher the cellular impact of 6p24.1 locus, we applied transcriptomic and epigenomic approaches on iPSC-derived endothelial cells to uncover potential genes regulated by the variants.

## Methods

### Study approvals and subject enrolment

This study was approved by the Local Ethics Committee of Nanyang Technological University Singapore Institutional Review Board (IRB18/09/02 and IRB-2020-09-011), National Healthcare Group (DSRB: 2013/00937) and Agency for Science, Technology and Research (IRB Reference 2020-096), Singapore. This research complies with the Helsinki Declaration. Written informed consent was obtained from each participant after the nature and possible consequences of the studies have been explained.

We leveraged on the ‘Singapore Coronary Artery Disease Genetics Study’ where CAD patients from the National University Hospital angiography center had been genotyped for rs6903956 alleles. CAD patients were diagnosed as non-ST-elevation myocardial infarction (NSTEMI) by angiography, while normal control subjects were clinically certified as healthy. This was to rule out normal controls who are asymptomatic but had undetected vessel blockage. Demographics of recruited subjects from whom samples were used for experimentation can be found in Supplemental Table S2.

### Peripheral blood mononuclear cell isolation and culture

For sample collection, 10 ml of fresh blood was collected from each subject in heparin vacutainers and processed in the laboratory within 6h. Upon centrifugation of blood using Ficoll-PaquePremium (GE Healthcare, catalog no.17-5442-03), a buffy coat layer containing peripheral blood mononuclear cells (PBMCs) were isolated. PBMC fractions were used for three purposes – (1) Cultivated in cell culture to derive induced pluripotent stem cells; (2) Analysis of circulating endothelial cells; (3) Genotyping (Supplemental Methods). For cryopreservation, PBMCs were resuspended in heat-inactivated FBS (Thermo Fisher Scientific Life Sciences, catalog no. 10082139) with 10% DMSO, frozen first at 80°C and then transferred to liquid nitrogen for storage.

### Sendai reprogramming of PBMCs to generate iPSCs

iPSC were generated from donor PBMCs by CytoTune-iPS Sendai Reprogramming (Thermo Fisher Scientific Life Sciences, catalog no. A16518). Please refer to Supplemental Methods for more details.

### Maintenance and characterization of induced pluripotent stem cells

iPSCs were grown on Matrigel-coated plates (Corning, catalog no. 354230) in mTeSR1 medium. Cells were passaged every 4 to 5 days using ReLeSR (StemCell Technologies, catalog no. 05872). We performed characterization of our iPSCs by immunofluorescence, karyotyping and teratoma formation assay. Please refer to Supplemental Methods for more details.

### Endothelial differentiation from iPSC lines

We followed our previously established differentiation protocols for lateral plate mesoderm derived endothelial cells by Cheung et al., 2012 and Narmada et al., 2016 [27, 28]. Please refer to Supplemental Methods for more details.

### Endothelial characterizations

Marker characterization of the endothelial cells from iPSCs were performed by flow cytometry and immunocytochemistry. Cells were washed and stained with primary antibodies, anti-human PECAM1 (CD31; Biolegend, catalog no. 102507) or anti-human ICAM1 (CD54; BioLegend, catalog no. 353111), in a staining buffer containing PBS with 2% heat-inactivated FBS, for 30 minutes in the dark. Following 30 minutes of antibody incubation at room temperature, stained cells were washed and resuspended in PBS containing 2% heat inactivated FBS. Flow cytometry data were collected on a BD LSRFortessa X-20 cell analyzer (Becton Dickinson) and analyzed using FlowJo v10.7.1 software (Becton Dickinson). Please refer to Supplemental Table S5 for details of primary antibodies.

Endothelial cells from iPSCs were functionally characterized by tube formation. 50 μl of Matrigel was added into each 96-well plate well and allowed to solidify for 30 min at 37 °C. Endothelial cells were seeded onto Matrigel at a cell density of 25,000 cells per well in EGM-2 and monitored every hour by light microscopy. Positive and negative controls were human coronary artery endothelial cells (HCAEC, ATCC, catalog no. PCS-100-020) and our derived iPSCs respectively. Phase-contrast images were acquired at 10× magnification using a Nikon Ti-E inverted microscope with MetaMorph version 7.8 (Molecular Devices, California, United States). We performed quantitative analysis of endothelial cord-like structures using Angiogenesis Analyzer on ImageJ [29].

ELISA of interleukin-8 was rendered on conditioned media of our iPSC-derived endothelial cells. Cells were seeded at 100,000 cells per well of 6-well plate in 2 ml of EGM-2. Culture media were conditioned for 72h and then harvested to undergo centrifugation at 13,000G for 10 min in order to remove cell debris. Supernatant fractions were collected for interleukin-8 ELISA (Abcam, catalog no. ab100575) according to manufacturer’s instructions. Normalization of IL-8 measurements was done against total cell protein of each well. Cells were washed with PBS before lysing with 350 ul RLT for total protein quantification using Pierce BCA Protein Assay Kit (Thermo Fisher Scientific Life Sciences, catalog no. 23227).

### RNA extraction and quantitative RT-PCR

Total RNA was isolated using RNeasy Plus Mini kit (Qiagen, catalog no. 74134) as per the manufacturer’s protocol, subsequently used to generate cDNA with LunaScript RT SuperMix Kit (New England Biolabs, catalog no. E3010S). Real-time PCR was performed using SYBR green gene expression assays (New England Biolabs, catalog no. M3003S) on a QuantStudio 6 instrument (Applied Biosystems). Gene expressions were normalized to endogenous GAPDH housekeeping gene. Please refer to Supplemental Table S5 for primer sequences.

### CRISPR-Cas9 genome editing at 6p24.1

Cas9 guide RNAs (gRNA) with predicted breaks were identified using a computational algorithm scoring system (https://benchling.com/). A non-homology end joining (NHEJ) double sgRNA-guided Cas9 system was used. Two gRNA duplex oligos were subcloned and ligated with pMIA3 plasmid [30] (Addgene #109399) using T4 DNA ligase (New England Biolabs, catalog no. M0202S). 1.5x10^6^ cells were pre-treated with 10 μM ROCKi (StemCell Technologies, catalog no. 72302) for an hour. Single cell suspension was prepared using accutase (StemCell Technologies, catalog no. 07922) and resuspended in 100ul of nucleofection solution and 10ug of pMIA3 plasmid containing the selected gRNA pairs. Cells were nucleofected using Amaxa4D nucleofector (Lonza, catalog no. AAF-1002B) and P3 primary kit (Lonza, catalog no. V4XP-3024) as per manufacturer’s instructions and plated out on matrigel coated 6-well plate using mTeSR with CloneR supplement. After 48 hours, cells were again pre-treated with ROCKi and a single cell suspension was prepared. Fluorescence activated cell sorting (FACS) was performed to enrich for targeted cells. RFP+ cells were sorted and plated onto 6-well plates containing mTeSR with CloneR, P/S and Gentamicin.

Colony formation was apparent from the individually sorted iPSCs 8 days after FACS. 24 single colonies then were manually picked into 12-well plates. Colonies were amplified and split with RelesR onto 6-well plates. Remaining cells were used for genomic DNA extraction and genotyping. Clones with successful CRISPR targeting were expanded. Refer Supplementary Figure S2a for gRNA sequences.

### ENCODE analysis

Histone marks in the 6p24 locus were queried from Roadmap epigenomics database by the ENCODE project. We downloaded the datasets from the ENCODE portal [31] (https://www.encodeproject.org/) with the following identifiers: ENCBS609ENC, ENCBS717AAA, ENCBS709TEL, ENCBS703ZDS, ENCBS899TTJ. H3K27ac and H3K4me3 histone marks were visualized on Integrated Genomics Viewer (IGV) using coordinates chr6:11,773,800-11,775,000 on hg38.

### ChIP-Seq analysis

Analysis of H3K27ac and H3K4me3 histone marks was conducted using coordinates chr10:44,297,249-44,312,248 on hg38 for visualization of weak promoter lying ∼2kb from fragment 4, and chr10:41,830,816-41,866,615 in hg38 for visualization of super-enhancer region. Individual bigWig (BW) files were visualized on IGV. ChIP-Seq human vascular endothelial cells was analyzed from GEO dataset (GSE131953).

### RNA-sequencing and analysis

Total RNA was isolated from the iPSC-derived endothelial cells as per manufacturer’s instructions (RNeasy Micro Kit, Qiagen, catalog no. 74004). PolyA library preparation and 150 base-pair paired end sequencing was performed by Novogene sequencing facility (Singapore) using an Illumina HiSeq sequencer. The average sequencing depth was 60 million reads. Reads were aligned using STAR aligner v2.4.1a to human genome (hg38) with default parameters. Read counts per gene were extracted from STAR output. Raw read counts were converted to log2-counts-per-million (logCPM) and the mean-variance relationship modelled with precision weights approach with voom transformation. For differential expression analysis, raw counts were normalized using edgeR R package v3.34.0 [32] using a trimmed mean of M values (TMM) and differentially expressed genes were identified using limma R package v3.46.0 [33]. Heatmaps were generated using pheatmap R package v1.16.0. Principal component analysis plots were generated with scatterplot3d R package v0.3-41. Gene ontology annotations [34] from Molecular Signatures Dataset [35] was used to perform Fast Gene Set Enrichment Analysis R package v1.18.0. The functional disease networks were generated through the use of IPA (QIAGEN Inc.) [36].

### Hi-C

Hi-C was performed using the Arima-HiC kit (Arima Genomics, San Diego), according to the manufacturer’s protocols up to purifying digested DNA. Purified DNA was then handed over to Integrated Genome Analytics Platform for steps up to library prep and sequencing. For creation and analysis of Hi-C contact maps, Raw fastq were first aligned to human reference genome hg38 in parallel using BWA-MEM (v0.7.15) with default parameters on each sample. Unmapped and abnormal chimeric reads were excluded [37]. Subsequently, duplicate reads and read pairs less than 2 kb apart were removed to avoid self-ligated fragments. The resulting .hic file contains filtered contact matrices, which were loaded into Juicebox for visualization.

For identification of toplogically associating domains (TADs), the resulting .hic matrix files (MAPQ > 30) were used as input for Arrowhead to identify TADs. Arrowhead ran automatically as part of Juicer pipeline post-processing. The following default parameters were set for Arrowhead: Resolution (r) = 5kb; Normalization (k) = Knight-Ruiz balancing (KR); Size of sliding window (m) = 2000.

For visualization of Hi-C output, Hi-C contact matrix are visualized using 3D genome browser and Juicebox software. Circular visualization of HUVEC (CC-2517) *trans* chromosomal interactions were visualized using Rondo [38].

### 3C-droplet digital PCR

3C libraries were prepared as previously described [39]. Briefly, 1 x 10^7^ cells were cross-linked with 2% formaldehyde for 10 minutes and quenched with 0.125mM glycine. Cells were lysed with 5mL cold lysis buffer (10 mM Tris-HCl, pH 7.5; 10 mM NaCl; 5 mM MgCl2; 0.1 mM EGTA; 1x complete protease inhibitor; 11836145001 Roche) for 10 minutes. Cell nuclei were pelleted and resuspended in 500uL 1.2x FastDigest Buffer (Thermo Fisher Scientific Life Sciences, catalog no. B64) with 0.3% SDS and incubated for 1 h at 37 °C. Triton X (final 2% v/v) was added and incubated for 1 h at 37 °C. Partially digested nuclei was incubated overnight at 37 °C with 400U of FastDigest *HindIII* (Thermo Fisher Scientific Life Sciences, catalog no. FD0504). To inhibit the restriction digestion, SDS was added (final 1.6% v/v) and incubated at 65 °C for 20-25 minutes, followed by Triton X (final 1% v/v) for an hour to sequester SDS. Digestion efficiencies were accessed using qPCR and only samples with the efficiency of restriction enzyme digestion above 60-70% were accepted for 3C analysis. For ligation of cross-linked and digested chromatin, samples were incubated at 16 °C for 4 h in the presence of 1.15x of T4 ligation buffer (New England Biolabs, catalog no. B0202S) and 100U T4 ligase. Proteinase K (Thermo Fisher Scientific Life Sciences, catalog no. EO0492) was added (300ug final) and samples were de-crosslinked overnight in 65 °C, followed by RNase (Thermo Fisher Scientific Life Sciences, catalog no. EN0531) digestion (300ug final). 3C samples were purified using phenol-chloroform extraction, and DNA concentration carefully determined via setting up qPCR reactions against reference genomic DNA of known concentration using “internal” primer sets amplifying GAPDH.

Probe-based ddPCR was performed following manufacturer’s protocol with reference to previously described 3C-digital PCR protocol [40]. Reactions were performed in total 24ul volume using 12ul of 2 x ddPCR Supermix for Probes (No dUTP) (Bio-Rad Laboratories, catalog no. 1863024), 900nM of target primer pairs, 250nM of probes and 400ng of 3C template DNA. To check for efficiency of probes and primers, a control primer priming the region upstream of the constant fragment was used as positive control. To exclude false positive results caused by non-specific background, ddH2O was used as negative control. Reactions were performed under universal cycling conditions: 95 °C for 10 min, followed by 45 cycles at 94 °C for 30s and 60 °C for 2 min and with final enzyme denaturation at 98 °C for 10 min. Signal was quantified using QX200 Droplet Reader (Bio-Rad Laboratories) and data analysis performed using Quantasoft Analysis Pro (v1.7). Only droplets above the minimum amplitude threshold determined from negative control wells containing no template DNA were counted as positive. The interaction frequency (= target copy number per 100ng sample) was calculated: 24 x copies/uL divided by 100 x amount of sample added to total reaction.

### Profiling of circulating endothelial cells in CAD patients

100µl of 1 million PBMCs were stained in the dark for 10 minutes at room temperature, followed by 20 minutes at 4°C on an analog tube rotator with antibodies (Supplemental Table S5: Resource Table). After incubation, cells were washed and resuspended in 200µl of PBS with 1% BSA for flow cytometry analysis. CECs were detected through the combined immunophenotypic profile of CD45-/CD31+/CD133-/DNA+. The number of CECs was expressed as cells per million of PBMCs. Flow Cytometry was performed using BD LSRFortessa X-20 and FACSDiva software (BD Biosciences) and data analyzed using FlowJo v10.7.1 software (Becton Dickinson). Each analysis included at least 30,000 events.

### RNA extraction and ddPCR in isolated CECs

Total RNA was isolated using RNeasy Plus Micro kit (Qiagen, catalog no. 74034) as per the manufacturer’s protocol, subsequently used to generate cDNA with LunaScript RT SuperMix Kit. Probe-based ddPCR was performed on QX200 Droplet Digital PCR System (ddPCR; Bio-Rad Laboratories) following manufacturer’s protocol. Absolute quantity of DNA per sample (copies/uL) was processed using QuantaSoft (v.1.0.596) and converted to copies/CEC according to amount of input sample. Please refer to Supplemental Table S5 for primer sequences.

### Statistical analysis

Statistical significance of differences between the cohorts were analyzed using GraphPad Prism version 9. Data normality was determined by Shapiro-Wilk test. Datasets with normal distributions were analyzed with unpaired Student’s two-tailed *t*-tests to compare two conditions; one- or two-way analysis of variance (ANOVA) followed by post-hoc Tukey for datasets with more than two conditions. Non-parametric Mann–Whitney *t*-test or Kruskal-Wallis test were used for non-normally distributed data. Application of these statistical methods to specific experiments is noted in the figure legends. A *p*-value of less than 0.05 were considered significant. Results are depicted as either mean ± standard deviation (SD) or mean ± standard error of mean (SEM). Power analysis was performed to compute the sample size (*n*) required to detect an effect that is true. A significance of 5% (α=0.05) and power of 80% (N=0.80) was used.

## Results

### Generation of endothelial cell models from coronary artery disease and normal subjects

In our subject recruitment, CAD patients were diagnosed as non-ST-elevation myocardial infarction (NSTEMI). We selected gender-, age- and ethnicity-matched CAD patients (homozygous for risk allele A at rs6903956) and normal subjects (homozygous for common allele G at rs6903956) for generation of their iPSCs (Supplemental Fig. S1a and Supplemental Table S2). CAD and normal iPSC lines (*n* = 4 from 2 donors/ group with 2 iPSC clones/ donor) were created through Sendai-based reprogramming of their respective peripheral blood mononuclear cell (PBMC) samples. The iPSC lines were characterized and showed tightly packed iPSC colonies expressing classical markers of pluripotency (Fig. 1a). Gold standard *in vivo* teratoma formation assay confirmed the presence of differentiated tissues characteristic of each germ layer from the histology sections of the teratoma (Fig. 1b), validating pluripotency of our derived iPSC lines.

Both CAD and normal iPSCs were also karyotypically normal (Supplemental Fig S1b). Endothelial differentiation was then performed on CAD and normal iPSCs using our established protocol [27, 28]. To better mimic coronary vasculature, iPSCs were first differentiated to obtain a lateral plate mesoderm population that is the precursor tissue for cell lineages of the heart (Fig. 1c). Subsequently, a high purity of PECAM1-expressing endothelial cells (> 98%) could be achieved after differentiation by chemically-defined conditions and fluorescence-activated cell sorting (Fig. 1c).

Endothelial cells derived from CAD and normal iPSCs (designated as CAD EC and normal EC respectively) expressed mature endothelial markers such as PECAM1, VWF and NOS3 (Fig. 1d), to comparable levels as the positive control, human coronary artery endothelial cells (HCAEC). Functionally, all iPSC-derived endothelial cell lines were capable of forming cord-like structures in tube formation assays (Supplemental Fig. S1c). We then determined if CAD EC could recapitulate some disease-relevant hallmarks. We found that CAD EC were more proinflammatory than normal EC by having greater ICAM1 expression (Fig. 1e), and secreting significantly higher amount of interleukin-8 (Fig. 1f), both of which are factors known to be implicated in atherosclerosis [41]. Next, we sought to establish the contribution of rs6903956 genotypes to endothelial phenotypes.

**Figure 1.**
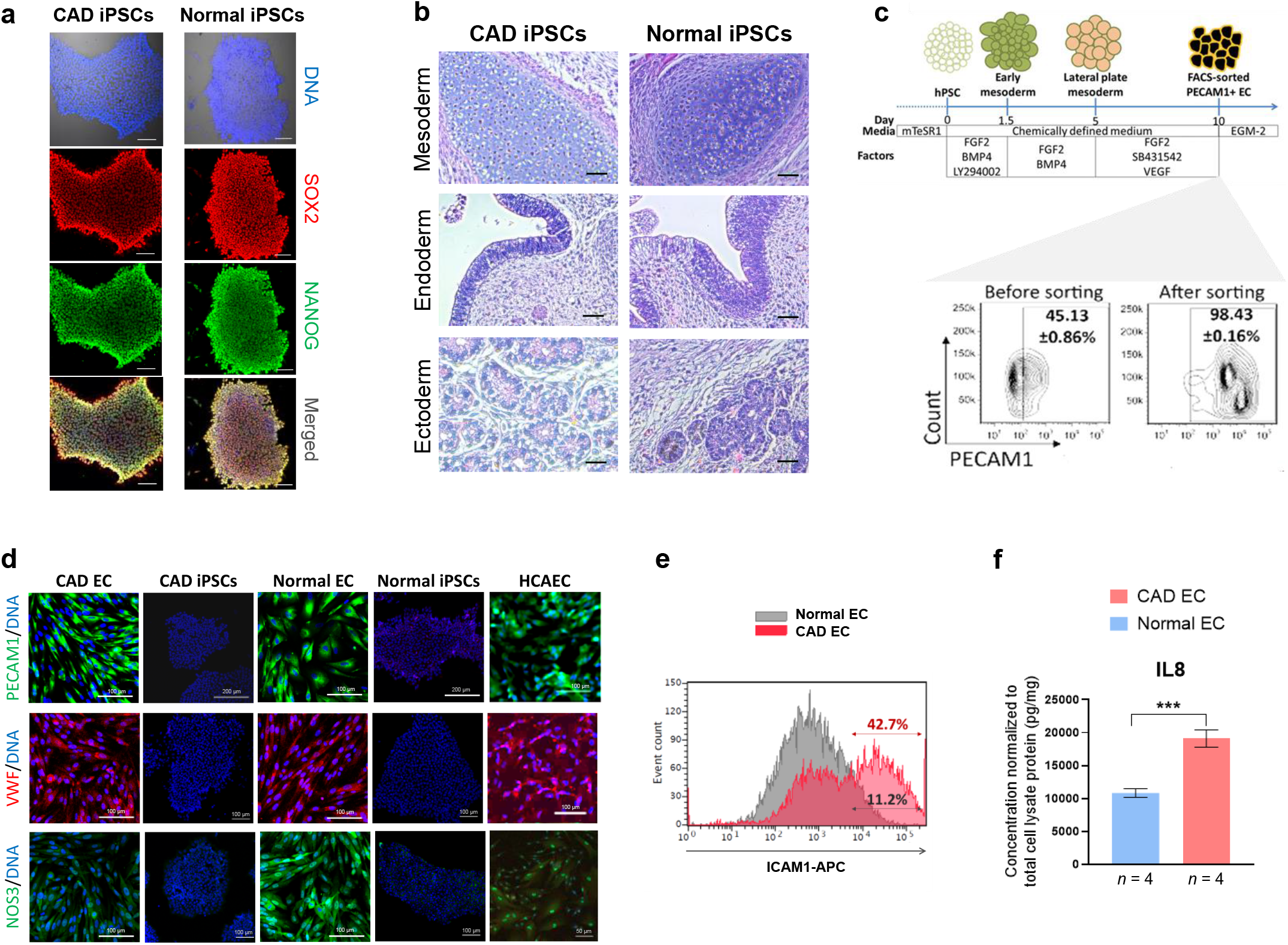
Derivation and characterization of induced pluripotent stem cells and endothelial cells from CAD patients and normal subjects. **(a)** Immunostaining of pluripotency markers, SOX2 and NANOG, on iPSC colonies (scale bar, 100 µm). **(b)** *In vivo* teratoma formation assay of derived iPSCs. H&E staining of teratomas for differentiated tissues of three distinct germ layers (scale bar, 25 µm). **(c)** Top: Stepwise protocol of iPSC differentiation toward endothelial lineage using chemically defined factors. Bottom: Representative flow cytometry plots of iPSC-derived endothelial cells for PECAM1 expression before and after FACS. **(d)** Immunostaining for mature endothelial markers on iPSC-derived endothelial cells from CAD patients and normal subjects (designated as CAD EC and normal EC respectively). Negative controls were CAD and normal iPSCs, while positive control was human coronary artery endothelial cells (HCAEC). **(e)** Flow cytometry characterization of proinflammatory marker ICAM1 on CAD EC and normal EC. **(f)** ELISA-based measurement of secreted interleukin-8 concentrations in culture media of CAD EC and normal EC, which were conditioned for 72 hours. Bar graphs showing means with S.D. (*n* = 4 from 2 donors with 2 iPSC lines/ donor), \*\*\**p*<0.001, two-tailed t-test.

### Effect of genome editing on cis gene expressions in iPSC-derived endothelial cells

*ADTRP* mRNA and protein expression have been detected in human endothelial cells [26, 42, 43]. To determine *ADTRP* expression in our iPSC-endothelial cells, we demonstrated that *ADTRP* was detected in both CAD EC and normal EC, as well as in the positive control HCAEC, but not in the iPSCs (Fig. 2a). We performed variant effect prediction [44] and rs6903956 showed up largely as an intronic variant as expected, with a minor possibility of being a potential regulatory region variant and nonsense-mediated mRNA decay (NMD) transcript variant (Fig. 2b). Diving deeper into the possible regulatory functions of rs6903956, we examined chromatin immunoprecipitation sequencing dataset [31] for the presence of active enhancer (H3K27ac) and promoter (H3K4me3) histone markers in relevant tissues and cells types. Spanning a 1,077bp region centered on rs6903956, there seemed to be negligible histone signals except for potential H3K4me3 marks in coronary artery (Fig. 2c). Although there were no obvious regulatory elements around rs6903956, it might be attributed to the lack of relevant endothelial cell models in the dataset.

If rs6903956 does not reside in regulatory regions, one of its potential impacts could involve long-range chromatin interactions with other genetic loci. Changes in chromatin interaction dynamics may affect the accessibility of regulatory regions distal to the SNP site, resulting in multifactorial cell-wide effects [45]. To understand tissue-specific chromatin architectures in our iPSC-derived endothelial model, we generated Hi-C genomic contact maps using CAD EC and normal EC by mapping chromatin contacts genome-wide at a 10-kb resolution (Fig. 2d). Using *in situ* Hi-C data, we obtained an average of 101 million long-range (≥ 20 kb) intra-chromosomal contacts after aligning and filtering 301 million Hi-C read pairs for each sample. Enhancers find their target genes within topologically associated domains (TADs), which are demarcated by interactome boundaries that restricts chromatin interactions within a spatially confined compartment in the genome [46]. As TADs are known to be strongly conserved across species and cell types, we compared CAD EC and normal EC Hi-C data with another well-characterized HUVEC line [47], which collectively reflected rather similar TAD characteristics (Fig. 2d).

To gain a better resolution of the effects of rs6903956, we performed genome editing to compensate for our limitation of not having iPSC lines from the same clinical phenotype with both risk and non-risk alleles. The minor allele A at rs6903956 has been involved as a CAD susceptibility SNP primarily in Chinese populations [8]. In analyzing nearby variants, a short tandem repeat variant, rs140361069, was found in weak linkage disequilibrium (LD) with rs6903956 in East Asian populations (R^2^ = 0.21), along with five other SNPs in strong LD (R^2^ > 0.8) (Fig. 4e). The ‘A’ risk allele at rs6903956 is correlated with deletion of (AAAT)3 at rs140361069 (Table 1). We postulated that the ethnicity-dependent effect of rs6903956 might be partially due to the lack of rs140361069 in LD with rs6903956 in Europeans and most of the other populations (Table 1). Hence, we generated isogenic iPSC lines with 63-89bp deletions on 6p24.1 containing rs6903956 and rs140361069 via Cas9-induced non-homologous end joining, with the use of a dual guide RNA (gRNA) targeting strategy [48] (Fig. 2f). To control for effects of CRISPR-Cas off-targeting, we designed two different pairs of gRNAs targeting the same region for both CAD and Normal iPSCs (Supplemental Fig. S2a). Guide RNAs were evaluated for DNA double strand break efficiencies in HEK293T cells using a disrupted GFP construct containing a 582bp *ADTRP* fragment (Supplemental Fig. S2b). In addition, we also generated unedited CAD and Normal iPSCs which were subjected to the same CRISPR-Cas9 nucleofection, but without a successful deletion at our target genomic region. Deletion of targeted genomic regions did not significantly impact iPSC growth and pluripotency marker expressions (TRA-1-60, OCT4, SOX2) (Supplemental Fig. S2c). These iPSC lines were further differentiated into endothelial cells. We henceforth designated WT CAD EC and WT Normal EC as the parental wild-type lines. Those with successful deletions of rs6903956 and rs140361069 were Δ CAD EC and Δ Normal EC, while those without successful deletions were UNΔ CAD EC and UNΔ Normal EC (Fig. 2f).

**Table 1:**
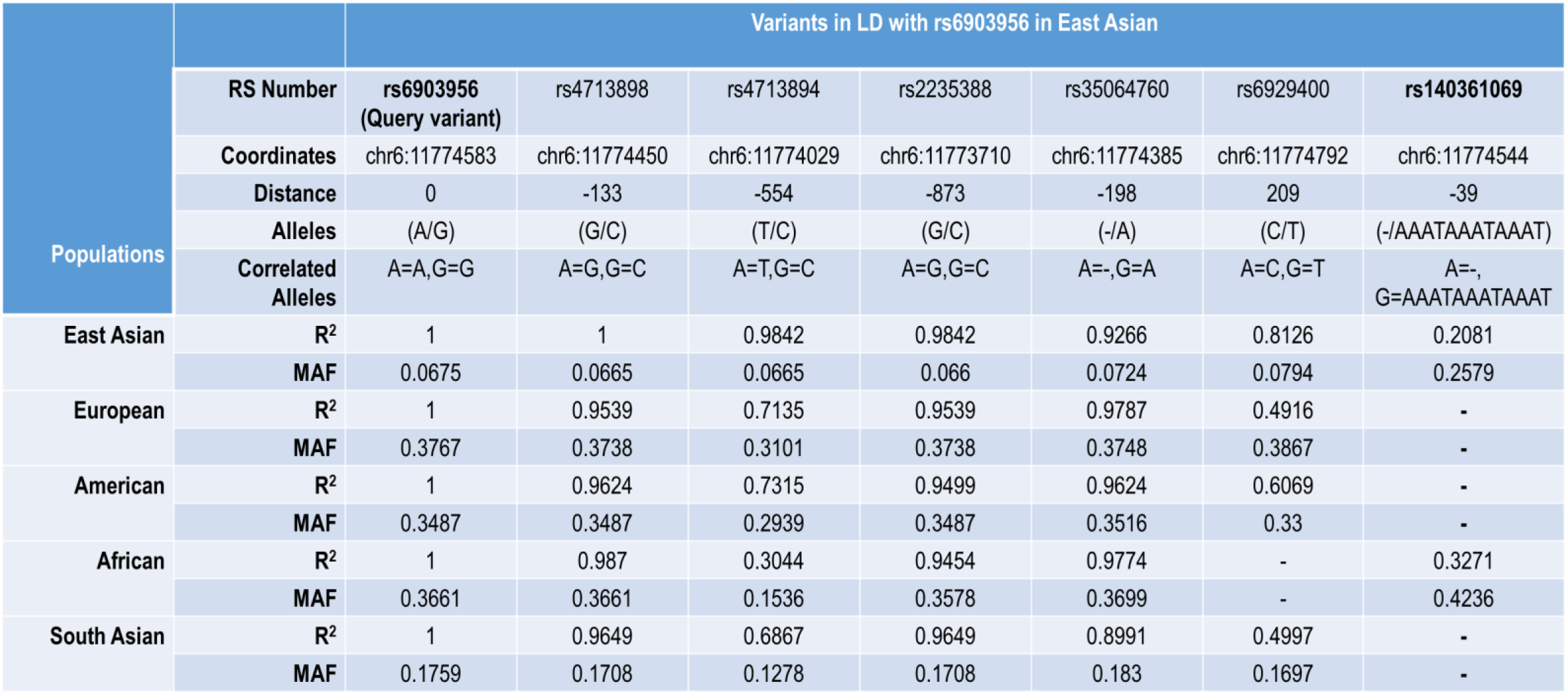
Variants in LD with the query variant rs6905956 in East Asian population are displayed. As a comparison, R2 value (cut-off > 0.2) and minor allele frequencies of those proxy variants were analyzed from other reference populations selected from the 1000 Genomes Project. Data have been retrieved from LDproxy Tool, part of the LDlink web-based applications for exploring LD in population groups [49].

| Populations | Variants in LD with rs6903956 in East Asian |  |  |  |  |  |  |  |
| --- | --- | --- | --- | --- | --- | --- | --- | --- |
|  | RS Number | rs6903956<br>(Query variant) | rs4713898 | rs4713894 | rs2235388 | rs35064760 | rs6929400 | rs140361069 |
|  | Coordinates | chr6:11774583 | chr6:11774450 | chr6:11774029 | chr6:11773710 | chr6:11774385 | chr6:11774792 | chr6:11774544 |
|  | Distance | 0 | -133 | -554 | -873 | -198 | 209 | -39 |
|  | Alleles | (A/G) | (G/C) | (T/C) | (G/C) | (-/A) | (C/T) | (-/AAATAAATAAAT) |
|  | Correlated Alleles | A=A,G=G | A=G,G=C | A=T,G=C | A=G,G=C | A=-,G=A | A=C,G=T | A=-,<br>G=AAATAAATAAAT |
| East Asian | R<sup>2</sup> | 1 | 1 | 0.9842 | 0.9842 | 0.9266 | 0.8126 | 0.2081 |
|  | MAF | 0.0675 | 0.0665 | 0.0665 | 0.066 | 0.0724 | 0.0794 | 0.2579 |
| European | R<sup>2</sup> | 1 | 0.9539 | 0.7135 | 0.9539 | 0.9787 | 0.4916 | - |
|  | MAF | 0.3767 | 0.3738 | 0.3101 | 0.3738 | 0.3748 | 0.3867 | - |
| American | R<sup>2</sup> | 1 | 0.9624 | 0.7315 | 0.9499 | 0.9624 | 0.6069 | - |
|  | MAF | 0.3487 | 0.3487 | 0.2939 | 0.3487 | 0.3516 | 0.33 | - |
| African | R<sup>2</sup> | 1 | 0.987 | 0.3044 | 0.9454 | 0.9774 | - | 0.3271 |
|  | MAF | 0.3661 | 0.3661 | 0.1536 | 0.3578 | 0.3699 | - | 0.4236 |
| South Asian | R<sup>2</sup> | 1 | 0.9649 | 0.6867 | 0.9649 | 0.8991 | 0.4997 | - |
|  | MAF | 0.1759 | 0.1708 | 0.1278 | 0.1708 | 0.183 | 0.1697 | - |

Our Hi-C contact maps earlier revealed rs6903956 to lie in TADs with neighboring genes *HIVEP1*, *EDN1* and *PHACTR1* (Fig. 2d). To identify if the deleted regions on 6p24.1 (Δ63-89bp) might affect expression of *cis* genes in the same TADs, we performed qRT-PCR with our edited and unedited endothelial cell lines. A previous study reported that presence of risk allele A at rs6903956 resulted in decreased mRNA expression of *ADTRP* in leukocytes [8]. However, no significant difference was observed for *ADTRP* and *HIVEP1* in our endothelial cell models (Fig. 2g). Interestingly, there was an upregulation of *EDN1* in Δ Normal EC only, suggesting a potential *cis*-regulatory effect due to the absence of non-risk G alleles at rs6903956. For *PHACTR1*, only significant baseline differences were observed between CAD and Normal ECs, instead of within their edited and unedited cell lines.

**Figure 2.**
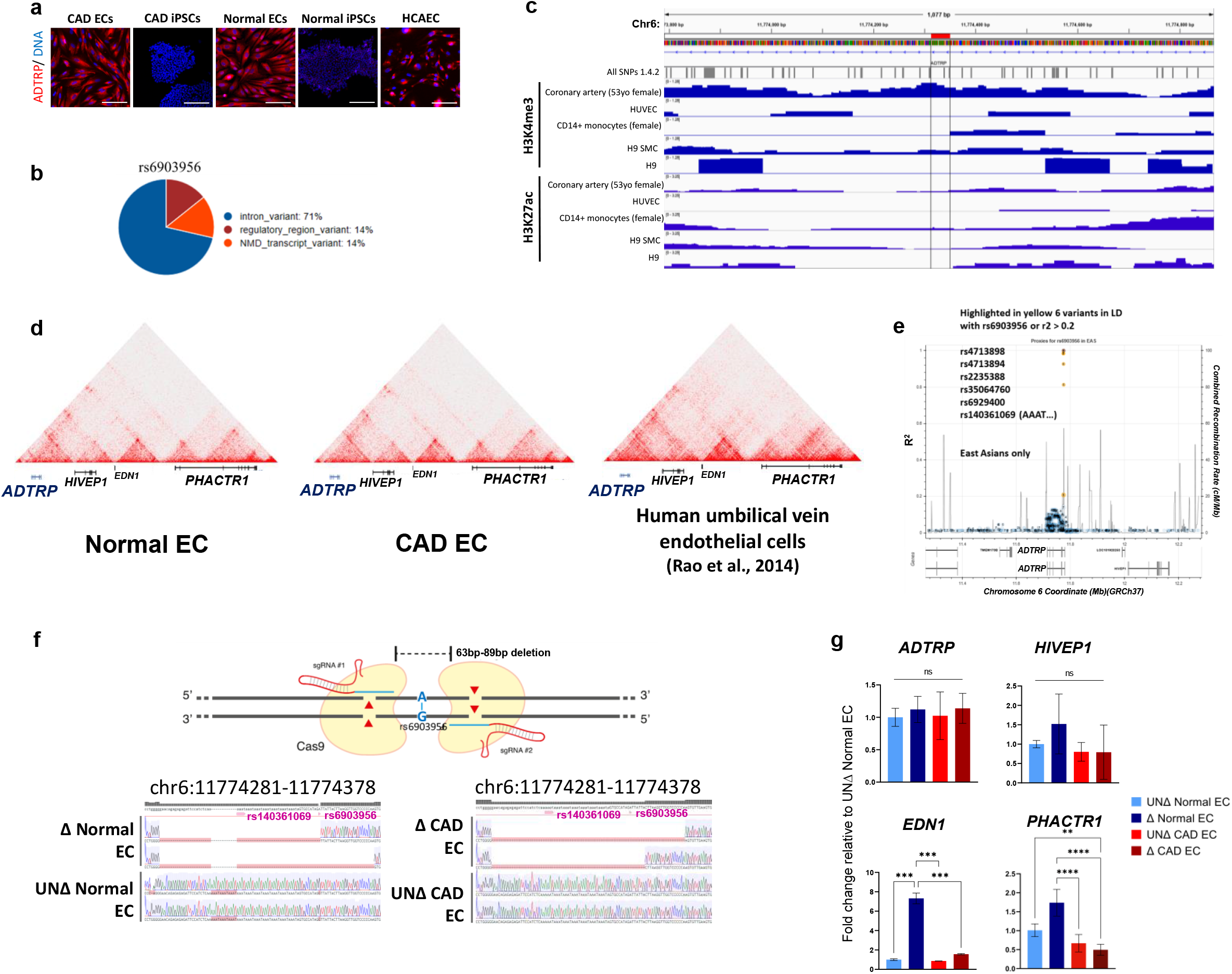
Investigation of rs6903956 *cis-*gene regulation through variant effect prediction and genome editing on iPSCs. **(a)** Immunostaining of ADTRP in CAD and normal iPSC-derived endothelial cells. Negative controls were iPSCs, while positive control was human coronary artery endothelial cells (HCAEC). Scale bar, 100 μm. **(b)** Annotation of rs6903956 by Ensembl Variant Effect Predictor **(c)** Visualization of H3K4me3 and H3K27Ac histone marks on hg38 chr10:44,297,249-44,312,248 based on chromatin immunoprecipitation sequencing (ChIP-seq) of various tissues/ cells. **(d)** Hi-C contact map visualization of 6p24.1 locus in Normal EC, CAD EC and HUVEC to reflect TADs involving rs6903956 in ADTRP and neighboring *cis* genes. **(e)** Regional association plot for ∼1kb region centered on query variant rs6903956 in CAD GWAS of East Asian Population from LDproxy tool [49]. Proxy variants with R^2^ > 0.2 were highlighted and identified as in linkage disequilibrium with rs6903956. **(f)** Top: Dual gRNA CRISPR-Cas9 deletion of 63-89bp regions flanking rs6903956 in WT CAD EC and WT Normal EC. Bottom: Sequencing chromatograms to validate CRISPR-Cas9 deletion in edited (Δ) versus unedited (UNΔ) endothelial cells. Two different sets of gRNAs were rendered on each iPSC-endothelial cell line to control for CRISPR-Cas9 off-targeting effects. **(g)** Quantitative RT-PCR of *ADTRP* and *cis* genes in isogenic edited and unedited CAD EC and Normal EC. Bar graphs showing means with S.D. (*n* = 4, from 2 donor cell lines with 2 biological replicates per cell line generated by 2 different guide RNA pairs), \*\*\**p*<0.001, one-way ANOVA.

### Genetically edited CAD endothelial cells reveal downregulation of molecular pathways relating to endothelial instability

To elucidate wider effects of the deleted region of 6p24.1 locus, RNA-sequencing was conducted on WT CAD EC with risk genotype (AA) and WT Normal EC with non-risk genotype (GG) (*n* = 3 biological replicates from 3 independent differentiation batches), as well as their edited (Δ) and unedited (UNΔ) counterparts (*n* = 6 from 2 cell lines generated by 2 different guide RNA pairs, 3 independent differentiation batches/ cell line). Principle component analyses demonstrated minimal variance between biological replicates (Supplemental Fig. S3a). There was also a high degree of similarity between different pairs of gRNAs targeting the same region, suggesting minimal off-targeting effect (Supplemental Fig. S3a). Differential expression analysis revealed 183 upregulated genes and 270 downregulated genes in Δ CAD EC compared to UNΔ CAD EC (Fig. 3a, top). Gene set enrichment analyses [35, 50] on these differentially expressed genes revealed downregulation of processes/ pathways relating to vasculature development, abnormal vascular physiology, endothelial cell migration and sprouting angiogenesis, etc (Fig. 3b). This might suggest that the deleted region on 6p24.1 containing AA risk genotype in Δ CAD EC had led to a less proliferative/ activated endothelial phenotype. On the other hand, there were substantially more differentially regulated genes between Δ Normal EC compared to UNΔ Normal EC (Fig. 3a, bottom). However, their gene set enrichment analyses did not reveal processes/ pathways that were directly indicative of perturbed vascular biology (Supplemental Fig. S3b). The deleted region on 6p24.1 containing GG non-risk genotype in Δ Normal EC might instead impact broadly on other fundamental biological processes. Therefore, we postulated that the rs6903956 G-to-A allele substitution could underlie a gain-of-function pathogenic role, as removal of AA risk genotype in Δ CAD EC, but not removal of GG non-risk genotype in Δ Normal EC, resulted in downregulation of endothelial activation mechanisms.

Next, we used a double-pronged approach to identify candidate effector genes due to the deleted region on 6p24.1. First, we performed ‘diseases and functions’ network analysis by Ingenuity Pathway Analysis on the differentially regulated genes between Δ CAD EC and UNΔ CAD EC. We found that ‘cardiovascular system development and function’ was among the most dysregulated network in Δ CAD EC (Supplemental Fig. 3c). Using the genes from this network, we derived a network interactome that showed predominately downregulated genes, with the CXC chemokine ligand 12 (*CXCL12*) being one of the central elements of this interactome (Fig. 3c). Secondly, based on parallel coordination plots of gene expression dynamics, we curated candidate genes whereby deletion of 6p24.1 containing AA risk genotype in Δ CAD EC could restore expressions to the levels found in WT normal EC harboring GG non-risk genotype (Fig. 3d). A heatmap visualization helped us appreciate the candidate genes that were normalized to the levels of WT normal EC upon removal of risk genotype in Δ CAD EC (Fig. 3e). Consistently, *CXCL12* surfaced as a potential candidate gene. *CXCL12* resides within chromosome 10q11.21 that have been reported as a risk locus for CAD susceptibility [6, 51]. Three *CXCL12* variants (rs266089, rs1065297, rs10793538) have been reported to show an association with CAD in the Chinese Han population [52]. We then genotyped our iPSC lines to ascertain if they had underlying *CXCL12* variants. All our iPSC lines harbored common alleles at those *CXCL12* SNPs with no known association with CAD, except for one of the normal iPSC lines with heterozygous ‘GA’ genotype at rs266089 that has been postulated to confer CAD risk (Supplemental Table S2). However, there has been no eQTL analysis to establish whether these *CXCL12* SNPs may result in change of *CXCL12* expression. We were motivated by the possibility of inter-chromosomal (*trans*) interaction of CAD susceptibility loci linking 6p24.1 and 10q11.21.

**Figure 3.**
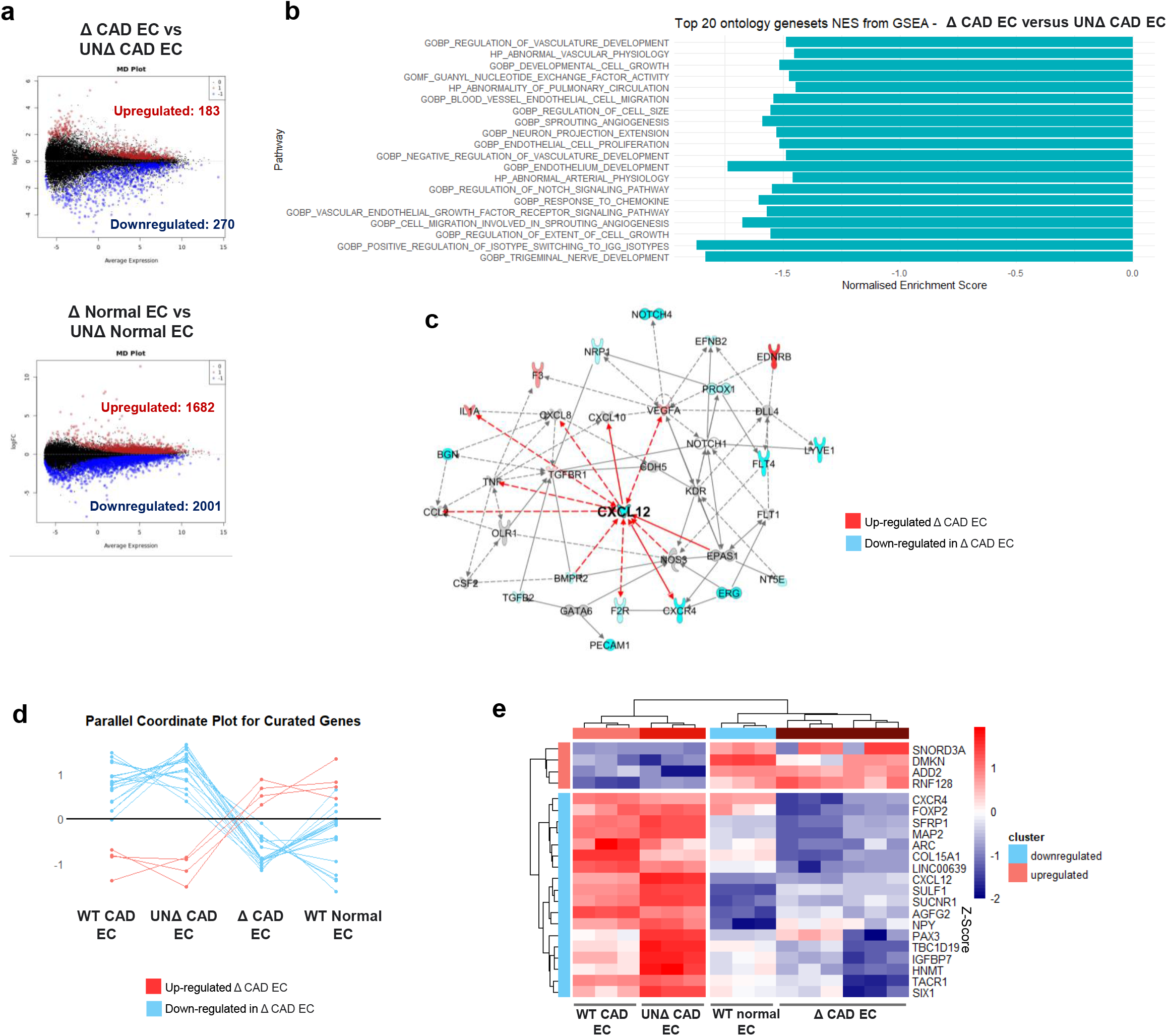
Transcriptomic analysis of isogenic edited and unedited iPSC-derived endothelial cells. **(a)** Mean-difference plots showing the log-fold change for all differentially expressed genes in Δ CAD EC versus UNΔ CAD EC (top), as well as in Δ Normal EC versus UNΔ Normal EC (bottom). **(b)** Top 20 ontology gene sets normalized enrichment scores from Gene Set Enrichment Analysis, ranked according to *p*-value. Log 2 fold change of genes differentially expressed in Δ CAD EC versus UNΔ CAD EC was used as input. **(c)** An interactome of differentially expressed genes in Δ CAD EC versus UNΔ CAD EC, arising from the ‘cardiovascular system development and function’ network based on Ingenuity Pathway Analysis. Genes indicated in red and blue were upregulated and downregulated in Δ CAD EC respectively. **(d)** Parallel coordinate plot for curated genes that showed gene expressions in Δ CAD EC ‘normalized’ to the levels of that in WT Normal EC. **(e)** Heatmap visualization of curated genes from (d).

### Trans interaction of 6p24.1 and 10q11.21 in endothelial cells

We performed data mining of Hi-C data from Rao et al. [47] to probe the chromatin landscape between 6p24.1 and 10q11.21 in HUVEC dataset. Strikingly, we observed that both chromosomal regions were closely intertwined in 3D space, probably due to the presence of a super enhancer region on 10q11.21, spanning around 30kb (Fig. 4a). Super enhancers are known to play key roles in organizing gene expression patterns that regulate cell identity [53]. To examine if such super enhancer activity was detected in other endothelial models, we analyzed a database of ChIP-seq on various human vascular endothelial cell lines [54]. Indeed, there was general increase in H3K27Ac enhancer marks at our observed super enhancer region on 10q11.21 in all vascular endothelial cell types (i.e. HCCaEC, HCoAEC, HaoEC, HPAEC, HENDC) except IMR90, a lung fibroblast cell line (Supplemental Fig. S4a).

While this super enhancer on 10q11.21 could have facilitated long-range physical interaction with 6p24.1, we asked if 6p24.1 covering rs6903956 could be in close genomic proximity with regulatory elements activating *CXCL12* transcription in CAD EC. Chromatin conformation capture (3C) was conducted on WT CAD EC with risk AA and WT normal EC with non-risk GG (*n* = 4 from 2 donor cell lines with 2 technical replicates per cell line) in order to probe interactions between a constant HindIII fragment covering rs6903956 (fixed interaction hotspot) and the entire *CXCL12* target genomic region covering 27 HindIII cutting sites (fragments 1-28) (Supplemental Fig. S4b). The complexity of the human genome makes it difficult to quantify true ligation frequencies due to other random competing interactions, especially since ligation frequency is generally low between two separate chromosomes [55, 56]. Furthermore, the inherent bias posed by 3C technique toward *cis* interactions due to the power-law decay model makes it only possible for the most frequent long-range interactions to be accurately quantified [56]. Therefore, instead of conventional TaqMan qPCR, we switched to droplet digital PCR, which delivered high precision of absolute ligation product copy numbers, without the numerous normalization controls that 3C-qPCR requires [40], allowing us to accurately quantify ligation frequency using 3C. To make up the large number of cells (10 million) required for 3C, we pooled five independently differentiated batches of iPSC-derived endothelial cells, at the same time controlling for batch variations. Our findings demonstrated frequent interactions between our control fragment containing rs6903956 with the entire *CXCL12* genomic region (Supplemental Fig. S4b). In particular, fragment 4 which lies within the 5’ untranslated region of *CXCL12* appeared to have the strongest interaction signals with the control fragment containing rs6903956 in WT CAD EC (Supplemental Fig. S4b). We repeated the experiment with prioritization on fragments 2-6 on WT CAD EC with risk alleles and WT normal EC with non-risk alleles. Similar results of interaction frequency peaking around fragment 4 in WT CAD EC were reproduced (Fig. 4b).

To understand if regions around fragment 4 could be functionally regulating *CXCL12* expression, we reanalyzed ChIP-seq dataset of vascular endothelial cells [54]. The data predicted weak and consistent promoter (H3K4me3) histone peaks lying on fragment 5, located 2kb from fragment 4 (Fig. 4c). Taken together, CAD background coupled with AA risk genotype at rs6903956 seemed to confer greater interaction frequency near a weak promoter on *CXCL12*.

**Figure 4.**
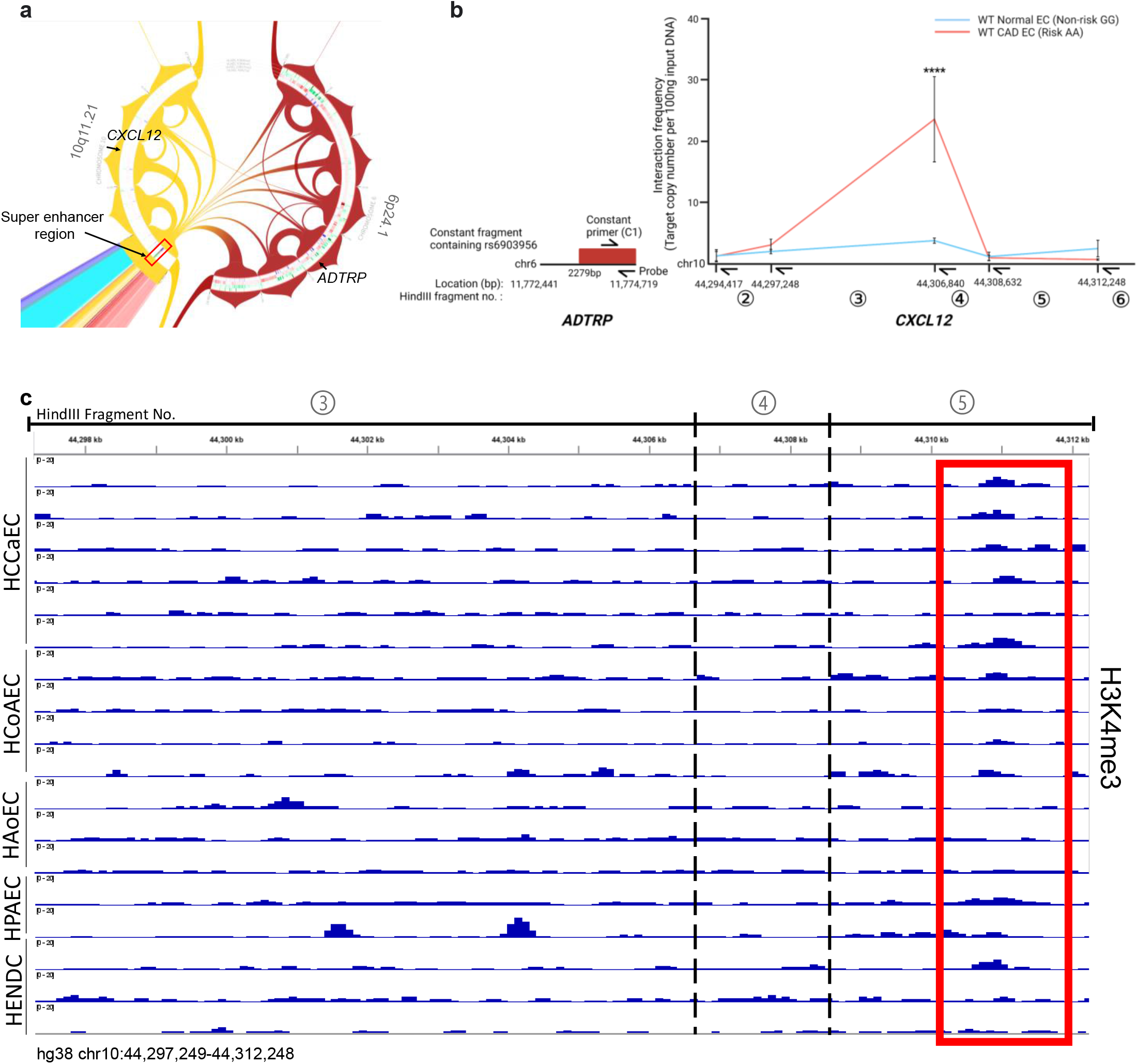
Frequent chromatin contacts between 6p24.1 and 10q11.21. **(a)** Chord diagram visualizing HUVEC dataset from Rao et al [47]. Hi-C inter-chromosomal loops between 6p24.1 and 10q11.21 on hg19. Only significant interactions (q-value<0.01) are shown. Presence of a super enhancer region on hg38: chr10:41,830,816-41,866,615 (boxed in red) where most of the chromatin contacts on 6p24.1 and 10q11.2 are anchored on, including *CXCL12* and *ADTRP*. **(b)** Chromatin conformation capture (3C)-droplet digital PCR of WT Normal EC and WT CAD EC. Anchoring on HindIII fragment harboring rs6903956 (constant fragment; 2279bp resolution), we probed for HindIII fragments 2-6 lying on 5’ untranslated region of *CXCL12*. Line graph showing means with S.E.M. (*n* = 4 from 2 donor cell lines with 2 technical replicates per cell line), \*\*\*\**p* ≤ 0.0001, two-way ANOVA comparing WT CAD EC with WT Normal EC for each individual fragment. **(c)** ChIP-Seq of H3K4me3 histone marks in vascular endothelial cells [54] on hg38 chr10:44,297,249-44,312,248: Common Carotid Artery ECs (HCoAEC, *n* = 4), Coronary Artery ECs (HCCaEC, *n* = 6), Human Aortic ECs (HAoEC, *n* = 3), Human Pulmonary Artery ECs (HPAEC, *n* = 2), Human Endocardiac Cells (HENDC, *n* = 3). Presence of a weak promoter ∼2kb downstream of fragment 4 was boxed in red.

### rs6903956 risk alleles are associated with vascular injury in patients with CAD

To test the effect of risk allele A at rs6903956, we measured circulating endothelial cells (CECs) as a biomarker for vascular injury [57], in the blood of CAD patients stratified by genotypes. CECs are damaged endothelial cells found in the peripheral blood, which have been detached from the blood vessel lining because of vascular injury [57–59]. We were mindful of our small sample size for iPSC-EC studies. Therefore, sample size derivation analysis was performed to ensure that validation experiments in CECs were adequately powered. We estimated fold effect size between risk genotypes AA/ AG and non-risk genotype GG, and their respective standard deviations based on flow cytometry detection of circulating endothelial cells in 8 samples per genotype group (Supplemental Table S3). The minimum number of samples needed to achieve statistical significance of p<0.05 with a power of 0.8 was 20 per genotype. With all CAD patients age- and gender-matched, we were able to achieve appropriate sample sizes in non-risk GG (*n* = 24) and risk AA/ AG (*n* = 31) genotypes (Supplemental Table S4). Remarkably, patients with risk genotypes AA/ AG had significantly higher number of CECs compared with patients with non-risk GG genotype (Fig. 5a). Our current results proved heightened levels of vascular injury associated with rs6903956 ‘A’ risk allele.

To validate variation of endothelial expression of *CXCL12* due to rs6903956, CECs were isolated by fluorescence-activated cell sorting and grouped according to the presence and absence of risk alleles (*n* = 27 ‘GA/AA’ and *n* = 20 ‘GG’). We performed droplet digital PCR to quantify *CXCL12* transcript copies due to the very low numbers of isolated CECs from the blood. We observed about 10 folds more *CXCL12* copy number per CEC in risk genotypes ‘AA/ AG’ compared with non-risk ‘GG’ genotype (Fig. 5b). These results strengthen the association between rs6903956 ‘A’ allele and higher *CXCL12* expression.

**Figure 5.**
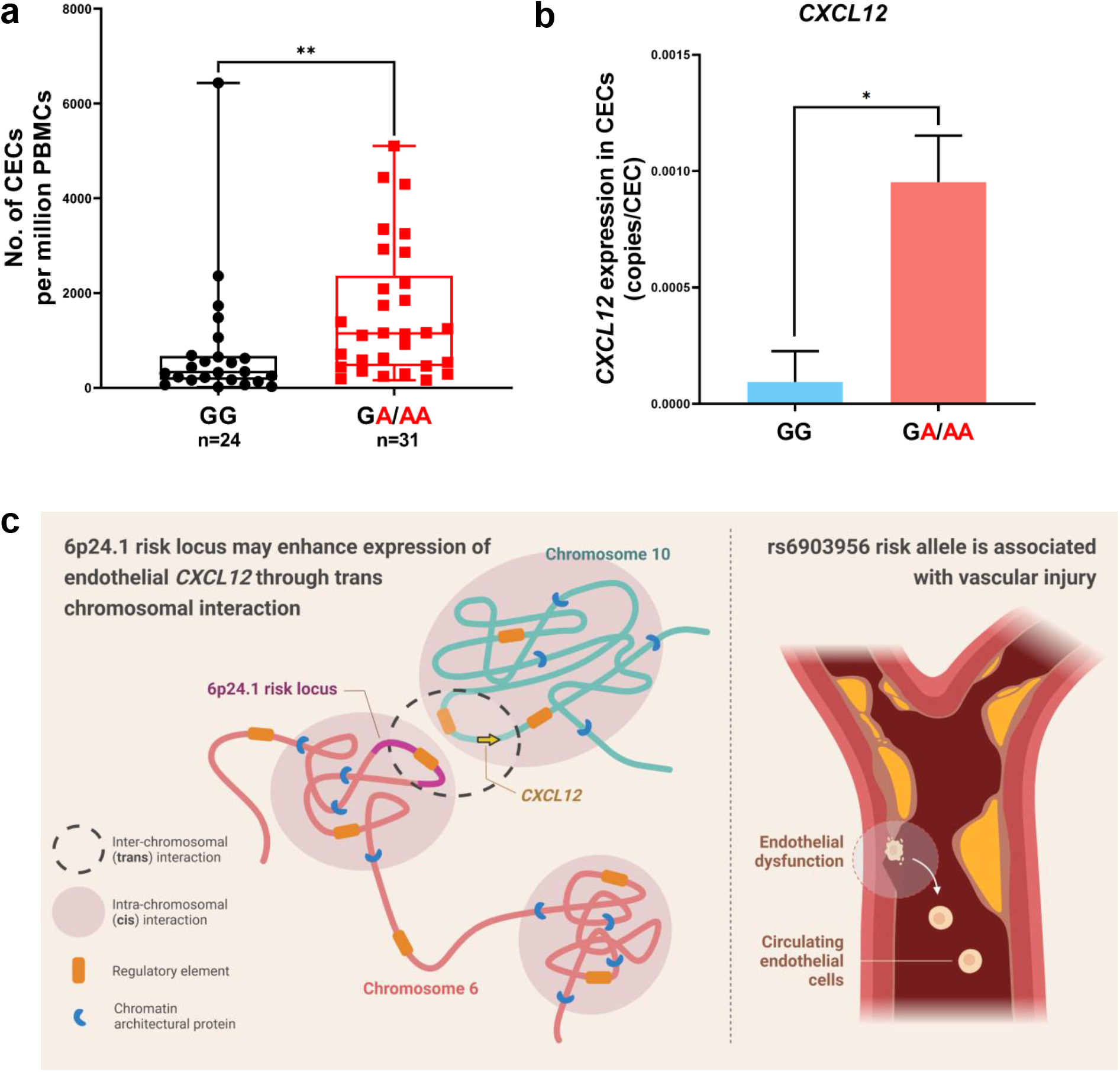
Profiling of circulating endothelial cells reveals association of rs6903956 risk allele with vascular injury. **(a)** Association between number of circulating endothelial cells (CECs) per million PBMCs in patient samples and their genotypes at rs6903956. Bar graphs showing means with S.D. (*n* = 31 ‘GA/AA’ and *n* = 24 ‘GG’), \*\**p* ≤ 0.01, Mann–Whitney *t*-test. **(b)** Droplet digital PCR of *CXCL12* transcript copies from isolated CECs pooled from ‘GG’ (*n* = 20) and ‘GA/AA’ (*n* = 27) patient samples. Bar graph showing means with S.D., \**p* ≤ 0.05, Mann–Whitney *t*-test. **(c)** Graphical summary of key findings.

## Discussion

Our genetic experimentation using patient iPSC-derived endothelial cells and data mining offers new insights into the molecular basis of susceptibility SNP rs6903956 in endothelial biology. We propose a mechanistic model in which trans-chromosomal impact on the atherosclerosis-implicated gene *CXCL12* could be one of the pathways through which 6p24.1 covering rs6903956 exerts its deleterious effects, as evidenced by elevated levels of vascular injury in patients associated with the A risk allele (Fig. 5b). Facilitated by CRISPR-Cas9 deletions (Δ63-89bp) on 6p24.1 including rs6903956 in our iPSC-derived endothelial cells, we demonstrated that presence of risk genotype AA at rs6903956 dysregulates vascular physiology transcriptional networks. On the other hand, removal of the non-risk GG genotype at rs6903956 had minimal effect on endothelial-specific pathways. Upregulation of *CXCL12* gene expression was related to increased inter-chromosomal interaction frequency of 6p24.1 with AA risk genotype near a putative *CXCL12* promoter region. Notably, a link between rs6903956 and vascular injury was validated by an association of risk allele A with higher numbers of damaged CECs detected.

SNP rs6903956 lies within the first intron of *ADTRP* that has been found to enhance the anticoagulant properties of endothelial cells in the presence of androgen [26]. *ADTRP*-deficient mouse and zebrafish models have exhibited multiple atherosclerotic hallmarks, such as leaky vessels, degradation of extracellular matrix and local inflammation [60]. ADTRP has also been found to specifically hydrolyze fatty acid esters of hydroxy fatty acids, which belong to a class of signaling lipids with anti-inflammatory properties [61, 62]. In human subjects, lower levels of plasma ADTRP have been reported in CAD patients compared with control subjects [14]. Nonetheless, there is still limited eQTL analysis performed on rs6903956 in endothelial system. Risk allele A at rs6903956 has been associated with decreased *ADTRP* mRNA expression in leukocytes [8]. More recently, the locus containing rs6903956 was postulated to act as an enhancer of *ADTRP*. Luo et al. transfected a 519bp region spanning rs6903956 and a 1513bp promoter/regulatory region of *ADTRP* into HeLa cells and found higher *ADTRP* promoter activity with preferential binding of transcription factor GATA2 to rs6903956 with common allele G [16]. They further showed in HUVECs that *GATA2* siRNA knock down resulted in lower ADTRP protein levels, but a direct enhancer-promoter interaction remained to be proven. As CAD pathogenesis also involves other cells such as smooth muscle cells and macrophage [3], cell-type specific effects of rs6903956 on *ADTRP* expression warrant further studies.

Causal non-coding SNPs can interfere with normal gene regulation by being regulatory loss- or gain-of-function mutations. On certain occasions, non-coding variants also have the potential to interfere with chromatin topological domain architecture by repositioning regulatory elements between domains or disrupting TAD boundaries, inducing ectopic gene activation and causing misexpression [63]. TADs form an intermediate level of chromatin organization by constraining interactions of *cis*-regulatory sequences with their cognate genes. We found previously that many cardiovascular disease-associated SNPs contribute to variation of gene expressions within the same TAD as the variants concerned [64]. Hi-C contact maps of our iPSC-derived endothelial cells revealed that *ADTRP* lies within the same TADs as neighboring genes *HIVEP1*, *EDN1* and *PHACTR1*. Alteration of *EDN1* expression was associated with non-risk homozygous G alleles. On the other hand, we demonstrated the feasibility of obtaining high-resolution *trans* interaction by coupling chromatin conformation capture (3C) with droplet digital PCR. Higher inter-chromosomal contact frequency was found between 6p24.1 containing rs6903956 and a 1,792bp region lying within 5’ untranslated region of *CXCL12*, which appeared to be a weak promoter. We note that rs6903956 is in linkage disequilibrium with rs140361069, a short tandem repeat which was missing in our CAD EC harboring homozygous A alleles at rs6903956. Sun et al. had shown that some disease-associated tandem repeats may be located with chromatin domain boundaries, affecting insulation of genomic neighborhoods [65]. It remains to be elucidated how G-to-A substitution at rs6903956 and/or missing tandem repeats at rs140361069 change chromatin dynamics, leading to elevated interaction frequency with a distal promoter site on *CXCL12*. No H3K27Ac enhancer histone marker has been detected at rs6903956 in vascular cells or tissues. Recent studies support the notion that enhancer activity might be the result of a synergistic action between multiple acetylation events at different histone sites, rather than the sole action of H3K27Ac [66]. Further investigation on other histone variants in endothelial cell system is necessary to substantiate the regulatory role of rs6903956 as an enhancer.

Our Hi-C data mining suggested there was an endothelial-specific super-enhancer on 10q11.21 that might be responsible for the regular contacts between 6p24.1 and 10q11.21. Super-enhancers have been thought to facilitate cell-identity specific regulatory response via the formation of phase separation bodies [67]. We find it fascinating that 6p24.1 and 10q11.21, both known loci harboring genetic polymorphisms associated with CAD, are closely interacting in 3D space. It will be worth investigating the potential global regulation of super-enhancers and how CAD risk loci on different chromosomes can mutually regulate gene expressions in chromatin space, opening the door for discovery of coregulated gene clusters via super-enhancer-mediated interchromosomal interaction.

In this study, rs6903956 was associated with increased *CXCL12* expression in endothelial cells. Serum CXCL12 levels are markedly increased in patients with cardiovascular diseases [68, 69], in which endothelial-derived CXCL12 is a key driver of atherosclerosis and one of the contributors to serum CXCL12 levels [70]. CXCL12 has previously been associated with a salutary effect on atherosclerotic plaque stability, possibly due to cell-specific atheroprotective effects and CXCL12’s role in recruitment of bone marrow-derived cells to sites of injury [71, 72]. In contrast, other studies have shown that inflammation mediated by CXCL12 and its receptors have long been linked to vascular injury, involving processes of monocyte differentiation and macrophage infiltration, aggravating pro-inflammatory responses on vascular endothelium [73]. CXCL12 is also involved in other pro-atherogenic processes, including hyperlipidemia and insulin resistance that may in turn affect endothelial integrity [74]. To validate our findings, we measured CECs as a biomarker to examine the functional consequence of rs6903956 on vascular injury in CAD patients, as CECs have been shown to be an indicator of vascular injury as they are shed into the circulation following vascular damage [75, 76]. CECs are valuable markers of vascular dysfunction in a variety of vascular disorders including myocardial infarction, acute ischemic stroke [77], atherosclerosis, vasculitis, coronary artery disease [78] etc. Interestingly, we detected a significantly higher number of damaged CECs in patients harboring risk alleles A at rs6903956 than those with homozygous non-risk alleles G. Furthermore, CECs isolated from patients with risk genotypes AA/ AG expressed higher levels of *CXCL12* gene expression compared with non-risk GG group.

There are limitations with our study. Firstly, it has been challenging to create single nucleotide switch from G-to-A at rs6903956. Hence, the findings based on edited iPSC-derived endothelial cells with small deletions (Δ63-89bp) on 6p24.1 may be confounded with unknown effects from uncharacterized regulatory elements within the deleted regions. Secondly, we have not validated if the 6p24.1 locus where rs6903956 resides has potential enhancer activity or converges on binding sites of chromosomal architectural proteins which may account for the differences of long-range chromatin interaction dynamics. Finally, we have shown a greater extent of endothelial damage associated with ‘A’ risk allele at rs6903956, but the direct consequence of higher *CXCL12* expression on vascular injury has not been demonstrated.

## Conclusions

Overall, our study demonstrates the advantage of gene editing in human iPSCs to uncover functional effects of a non-coding disease-associated SNP relevant to endothelial biology in patients with CAD. Implementation of genetic testing is cost-effective for triaging patients based on individual genetic susceptibilities. Our findings of rs6903956-associated pathways may improve risk stratification and guide the design of therapeutic trials aiming at mitigating CAD. This will serve as a proof-of-principle that patient-derived iPSCs can offer valuable insights into precision medicine approaches of genotype-guided therapies.

## Nonstandard Abbreviations and Acronyms

ADTRP: Androgen-dependent tissue factor pathway inhibitor regulating protein
CAD: Coronary Artery Disease
CRISPR: Clustered regularly interspaced short palindromic repeats
ChIP-Seq: Chromatin Immunoprecipitation Sequencing
CECs: Circulating endothelial cells
EGM-2: Endothelial growth medium-2
eQTL: Expression quantitative trait loci
FACS: Fluorescence activated cell sorting
GWAS: Genome-wide association Studies
gRNA: guide RNA
HUVEC: Human umbilical vein endothelial cell
iPSC: Induced pluripotent stem cell
NHEJ: Non-homologous end joining
PBMC: Human peripheral blood mononuclear cell
PCR: Polymerase chain reaction
SNP: Single nucleotide polymorphism
TAD: Topologically associating domain
3C: Chromosome conformation capture

## Acknowledgements

We thank all patients and healthy donors who have participated in this study. Special thanks to the team at National University Hospital for coordinating clinical sample collection, Ms Kee Bee Leng for conducting interview with each participant, Mr Tan Wei Heng, Ms Konstanze Tan and Mr Marcus Teo for experimental assistance, Asst Prof Melissa Fullwood and Dr Wilson Lek Wen Tan for discussion on ChIA-PET and Hi-C visualizations. We thank Dr. Kao Shih Ling for initial assistance with recruiting healthy donors for human iPSC generation.

## Authors’ contributions

Conceptualization: CC

Data curation: KYT, KXW, FWJC, NMQP, BCN, HHL

Formal analysis: KYT, KXW, FWJC, CKH, MYYC, CC

Investigation: KYT, KXW, FWJC, CC

Methodology: KYT, KXW, MIA, FWJC, BCN, ST, AFL, MML, EGYC, HHL, AKKT, JNF, RSYF, CC

Resources: CC, SLK, MYYC, CKH

Validation: KYT, KXW, FWJC

Visualization: KYT, KXW, FWJC, CC

Funding acquisition: CC, MYYC, CKH, RSYF, AKKT

Project administration: CC, MYYC, CKH

Supervision: CC, MYYC, CKH, RSYF, JNF, AKKT

Writing – original draft: KYT, CC

Writing – review & editing: All authors

All authors approved the submitted manuscript version.

## Data Availability

The authors declare that all data supporting the findings of this study are available within the paper and supplemental materials. Specifically, RNA-sequencing dataset that support the findings of this study are available in the GEO repository (https://www.ncbi.nlm.nih.gov/geo/query/acc.cgi?acc=GSE195656).

## Research Materials Availability

Some materials used in this study are commercially procured. There are restrictions to the availability of induce pluripotent stem cell lines derived from human patients and normal donors due to ethics considerations for use of these materials within the current scope of study. Requests can be made to the corresponding author as we will explore use of materials subject to new ethics approval and research collaboration agreement (including material transfer).

