## Supplemental for "*Trans*-interaction of risk loci 6p24.1 and 10q11.21 is associated with endothelial damage in coronary artery disease"

### Supplemental Information

#### Supplemental Figures and Legends

##### Supplemental Figure S1

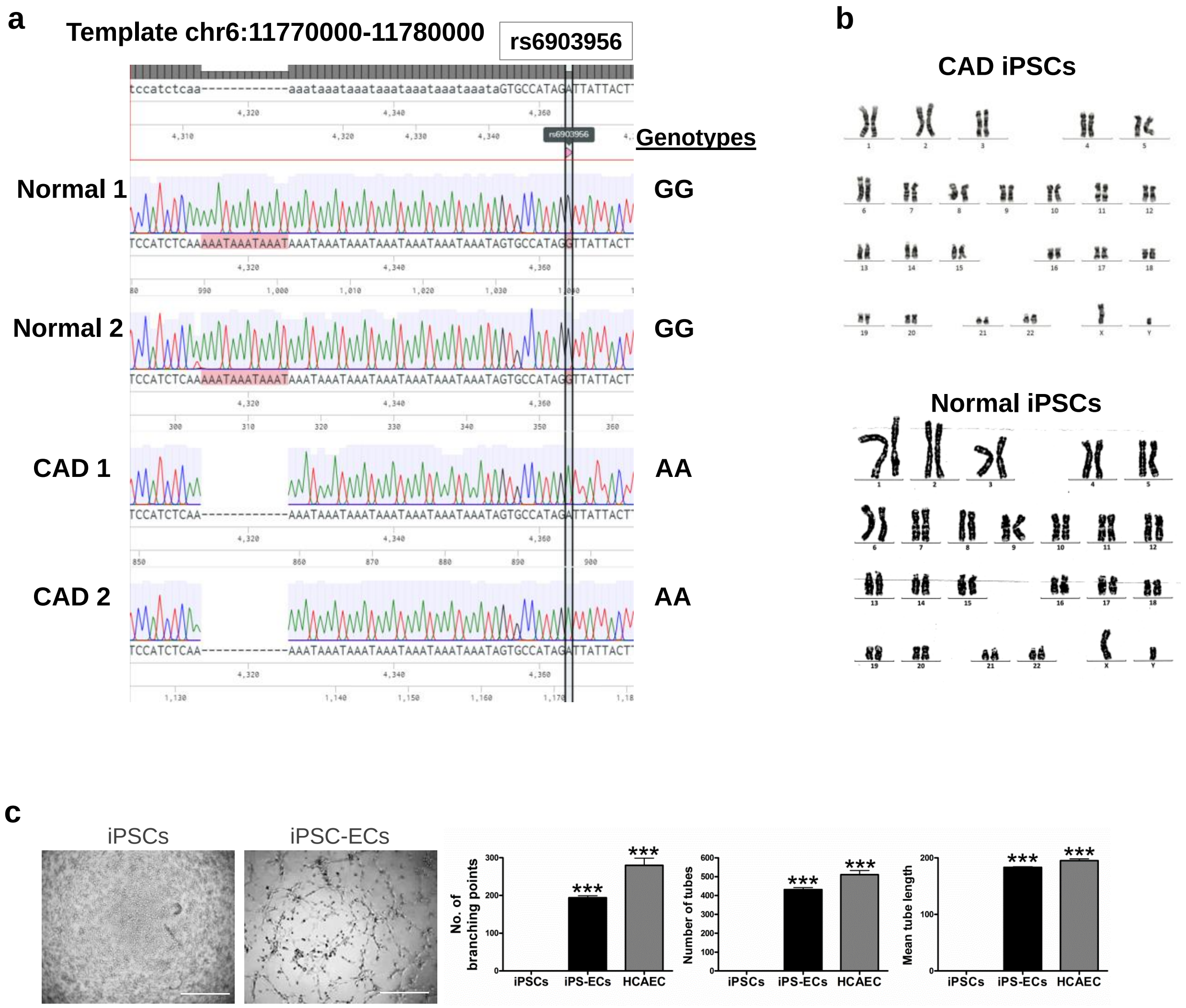


**Supplemental Figure S1: Characterization of induced pluripotent stem cells and endothelial cells from CAD patients and normal subjects. (a)** Genotyping of PBMC samples from 2 CAD patients and 2 normal subjects at 6p24.1 locus to ascertain genotypes at SNP rs6903956. **(b)** Karyotyping of CAD iPSC and normal iPSC to establish genetic stability of cell lines based on chromosome numbers and structural features. **(c)** Left: Representative images of tube formation after 2 hours by iPSCs and iPSC-derived endothelial cells (scale bar, 200 μm). Right: Image quantification of tube formation attributes of endothelial cord-like structures. Negative control was iPSCs, while positive control was human coronary artery endothelial cells (HCAEC). Bar graphs showing means ± s.e.m. ($n$ = 3 biological replicates), ***$p$<0.001, one-way ANOVA.

#### Supplemental Figure S2

a

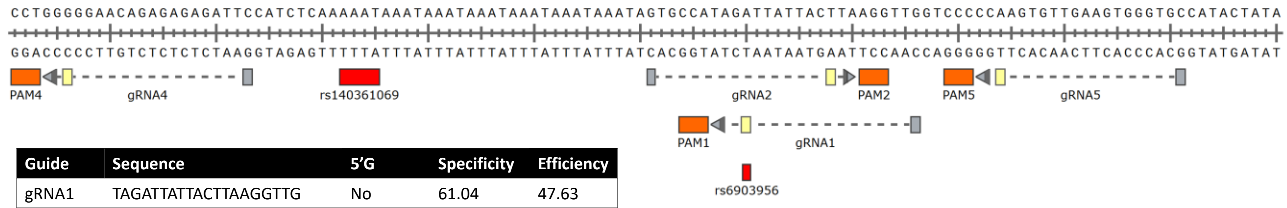


| Guide | Sequence | 5'G | Specificity | Efficiency |
|---|---|---|---|---|
| gRNA1 | TAGATTATTACTTAAGGTTG | No | 61.04 | 47.63 |
| gRNA2 | GTGCCATAGATTATTACTTA | Yes | 65.11 | 42.73 |
| gRNA4 | GGGGGAACAGAGAGAGATTC | Yes | 54.53 | 53.74 |
| gRNA5 | CCAAGTGTTGAAGTGGGTGC | Yes | 71.61 | 70.20 |

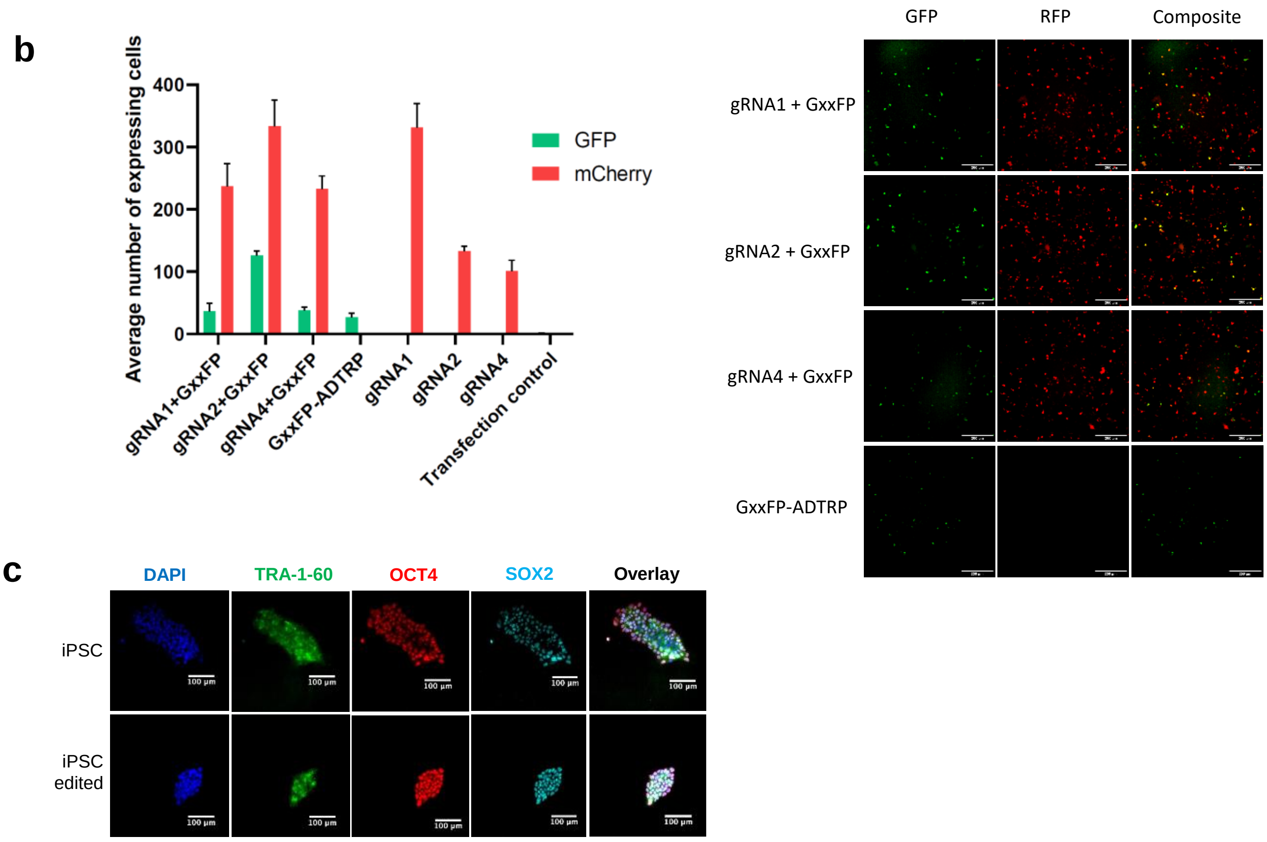


**Supplementary Figure S2. Genome editing strategy. (a)** Top: Location of gRNA sequences flanking rs6903956 and rs140361069 (chr6:11,714,280-11,714,400). Below: Table summarizing individual gRNA sequences with specificity and efficiency scores ranked by algorithm developed by Doench et al(Doench, Fusi et al. 2016). **(b)** pMIA3 containing a respective gRNA was co-transfected with pCAG-EGxxFP plasmid(Mashiko, Fujihara et al. 2013) (Addgene plasmid # 50716), a gift from Masahito Ikawa, disrupted by 582bp *ADTRP* region. gRNAs 1, 2 and 4 were evaluated for targeting efficiencies by quantifying the average number of expressing cells. pMIA3 plasmids without cotransfection of pCAG-EGxxFP was used as control. Inactive pMIA3 plasmid was used as a negative control. Bar graph showing means with S.D. (*n* = 3 independent transfections). **(c)** Immunostaining for pluripotency markers on iPSCs and genetically edited iPSCs (scale bar, 100 µm).

#### Supplemental Figure S3

a

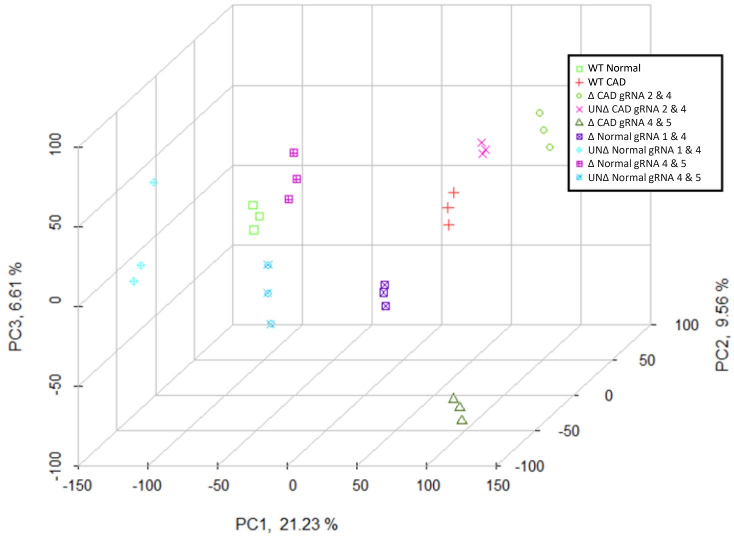


b

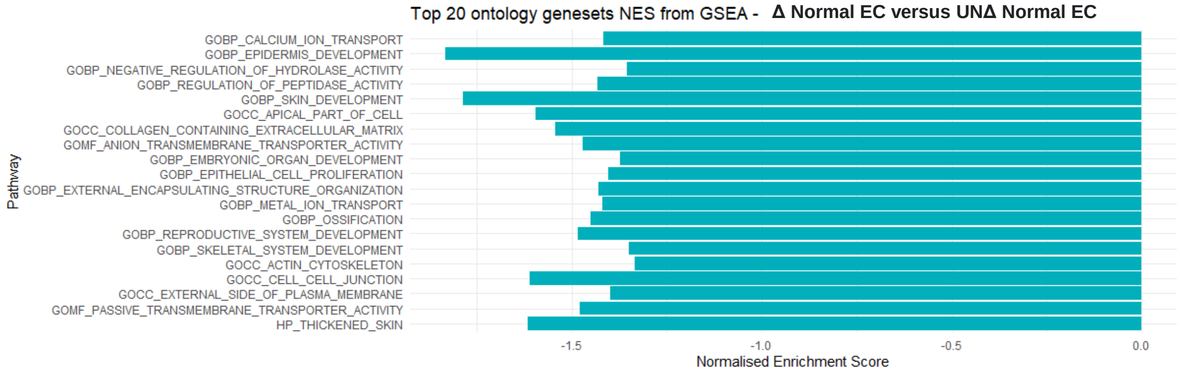


c

| ID | Molecules in Network | Score | Focus Molecules | Top Diseases and Functions |
|---|---|---|---|---|
| 1 | *BGN,BMPR2,CCL2,CDH5,CSF2,CXCL10,CXCL8,CXCR4,DLL4,EDNRB,EFNB2,ERG,F2R,F3,FLT1,FLT4,GATA6,IL1A,KDR,LYVE1,NOS3,NOTCH1,NOTCH4,NRP1,NT5E,PECAM1,PROX1,TGFB2,TNF,VEGFA* | 8 | 18 | Cardiovascular System Development and Function<br>Cellular Development<br>Cellular Function and Maintenance |
| 2 | *CXCL12,EPAS1* | 0 | 1 | Cardiovascular System Development and Function<br>Organ Morphology<br>Organismal Development |
| 3 | *ANGPT1,TEK* | 0 | 1 | Cellular Development<br>Embryonic Development<br>Tissue Development |
| 4 | *OLR1,TGFBR1* | 0 | 1 | Cardiovascular System Development and Function<br>Cell Death and Survival<br>Cell Morphology |

**Supplemental figure S3. Transcriptomic analysis of isogenic edited and unedited iPSC-derived endothelial cells (a)** 3D-PCA plot of wild-type iPSC-derived endothelial cells (*n* = 3 biological replicates from 3 independent differentiation batches), as well as edited and unedited endothelial cells (*n* = 6 from 2 cell lines generated by 2 different guide RNA pairs, 3 independent differentiation batches/ cell line). **(b)** Top 20 ontology gene sets normalized enrichment scores from Gene Set Enrichment Analysis, ranked according to *p*-value. Log 2 fold change of genes differentially expressed in Δ Normal EC versus UNΔ

Normal EC were used as input. **(c)** ‘Diseases and functions’ network analysis by Ingenuity Pathway Analysis for differentially expressed genes in Δ CAD EC versus UNΔ CAD EC.

#### Supplemental Figure S4

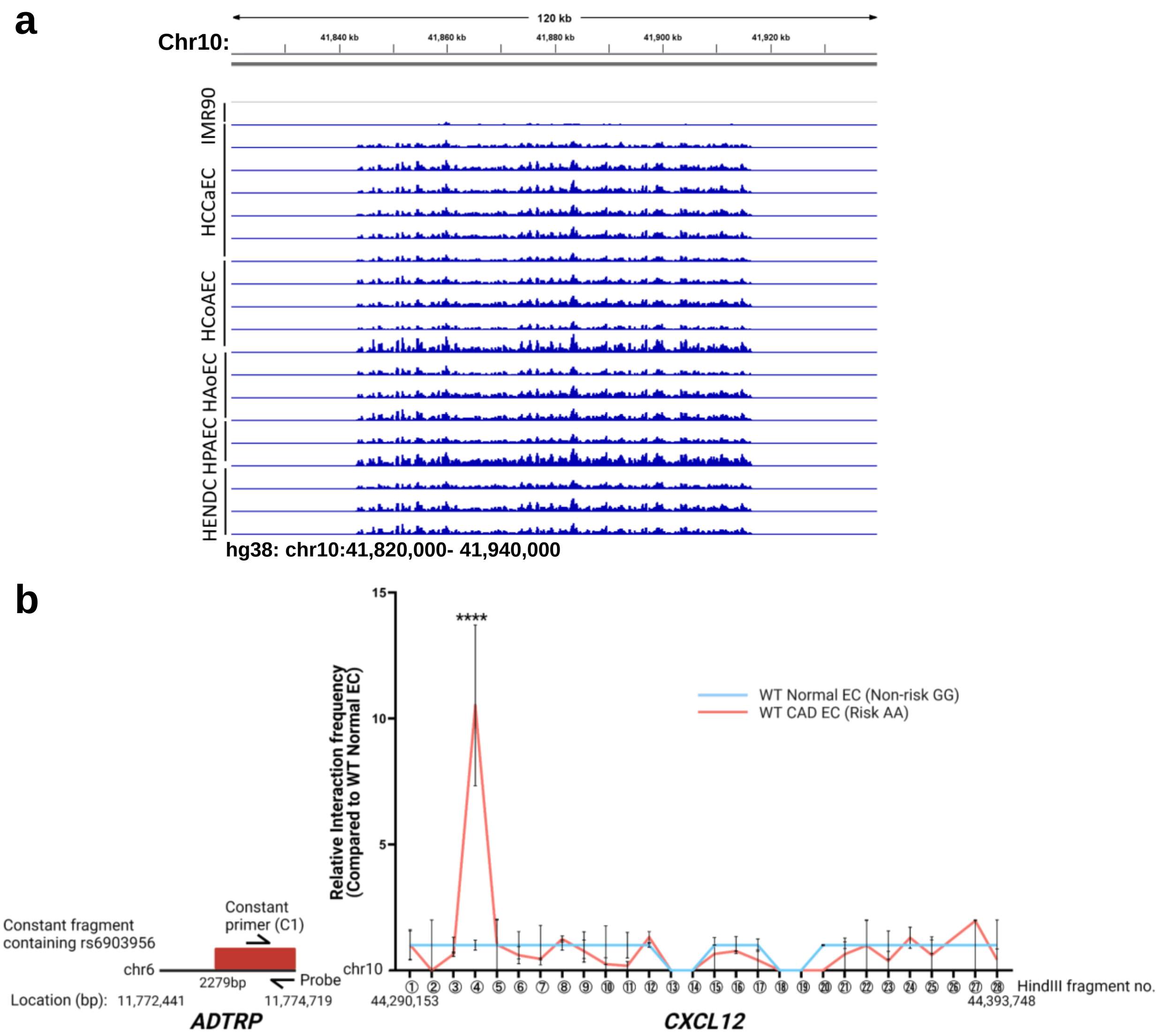


**Supplemental figure S4. Chromatin landscape of 6p24.1 and 10q11.21 (a)** ChIP-Seq of H3K27Ac histone marks in vascular endothelial cells(Nakato, Wada et al. 2019). Presence of an endothelial-specific super enhancer region on hg38: chr10:41,830,816-41,866,615. Normal lung fibroblast cell line (IMR90) was used as a non-vascular control. **(b)** Chromatin conformation capture (3C)-digital PCR of WT Normal EC and WT CAD EC. Anchoring on HindIII fragment harboring rs6903956 (constant fragment; 2279bp resolution), we probed for HindIII fragments for the entire *CXCL12* gene locus. Data represent fold changes against WT Normal EC. Line graph showing means with S.E.M. (*n* = 4 from 2 donor cell lines with 2 technical replicates/ cell line), ****$p \leq 0.0001$, two-way ANOVA comparing WT CAD EC with WT Normal EC for each individual fragment.

#### Supplemental Tables

**Supplemental Table S1**: Odds ratios of rs6903956 from different patient cohorts. Adjusted p-values and odds ratios were obtained after adjustment for covariates.

| Cohort description | Associated risk | rs6903956 | Adjusted Odds Ratio (95% CI), *p*-value | Minor Allele Frequency (A) (%) |
|---|---|---|---|---|
| Chinese Han population of 3,470 cases and 4,583 controls(Wang, Xu et al. 2011). | Coronary Artery Disease | G vs. A | 1.65 (1.44-1.90),<br>p=2.55 × 10−13<br>in the combined populations | Cases = 10<br>Controls = 7 |
| A cohort of 1,075 Chinese patients who underwent coronary arteriography(Guo, Gu et al. 2012). | Angiographical characteristics of coronary atherosclerosis risk | GG vs. AG | 1.444 (1.036-2.013),<br>p=0.030 | 9.07 |
| | | GG vs. AA | 5.896 (1.299-26.750),<br>p=0.021 | |
| 645 CAD subjects were recruited from patients undergoing coronary artery bypass graft, comprising of 406 Chinese, 122 Malays and 117 Indians(Tayebi, Ke et al. 2013).<br><br>755 controls were consecutive subjects attending their routine medical examinations. The ethnic composition for controls was 421 Chinese, 118 Malays and 216 Indians. | Coronary Artery Disease | GG vs. AG | Chinese=2.26 (1.09-4.6),<br>p=0.028<br>Malays=2.64 (0.6–11.59),<br>p=0.19<br>Indians=1.66 (0.7–3.4),<br>p=0.17 | Cases = 11.6<br>Controls = 7.4 |
| | | GG vs. AA | Chinese=1.26 (0.06–25.3), p=0.87<br>Malays=nil<br>Indians=1.11 (0.1–8.7),<br>p= 0.91 | |
| 1,536 consecutive autopsy cases of elderly Japanese patients(Dechamethakun, Ikeda et al. 2014). | Coronary stenosis index | G vs. A | 1.839 (1.172-2.886),<br>p=0.008 | Cases = 4.4<br>Controls = 7.3 |

**Supplemental Table S2:** Demographics of CAD patients and normal subjects from whom samples were used for development of iPSC-based endothelial models. At least two iPSC clones per subject were characterized for downstream experiments.

| Generation of iPSC-based endothelial cell models | | | | | | | | |
|---|---|---|---|---|---|---|---|---|
| Disease status | Diagnosis | Ethnicity | Gender | Age | CAD-associated SNP rs6903056 on 6p24.1 (risk allele: A) | rs1065297 on 10q11.21 | rs10793538 on 10q11.21 | rs266089 on 10q11.21 |
| | | | | | Related to Fig. 1 | Related to Fig. 3 | | |
| CAD | NSTEMI | Chinese | M | 68 | AA | AA | AA | GG |
| | NSTEMI | Chinese | M | 63 | AA | AA | AA | GG |
| Normal | Normal | Chinese | M | 58 | GG | AA | AA | GA |
| | Normal | Chinese | M | 56 | GG | AA | AA | GG |

**Supplemental Table S3**: Sample size derivation for circulating endothelial cell profiling based on power analysis.

| | | |
|---|---|---|
| Confidence Interval (2-sided): | 95% | |
| Power: | 80% | |
| Ratio of sample size (non-risk/ risk): | 1 | |
| Genotypes | Risk **AA / AG** | Non-risk **GG** |
| No. of circulating endothelial cells per million PBMCs | 1391.27 | 684.71 |
| | 242.39 | 1062.24 |
| | 290.37 | 226.44 |
| | 1160.39 | 26.75 |
| | 3352.09 | 652.83 |
| | 586.45 | 340.77 |
| | 292.02 | 133.61 |
| | 2205.40 | 557.05 |
| Mean | 1190.05 | 460.55 |
| Standard deviation | 1110.017 | 342.8438 |
| Sample size of AA/AG | **20** | |
| Sample size of GG | **20** | |
| Total sample size | **40** | |

*Results from OpenEpi, Version 3, open source calculator (https://www.openepi.com/SampleSize/SSMean.htm)*

**Supplemental Table S4:** Demographics of CAD patients from whom samples were used for analysis of patient circulating endothelial cells.

| Analysis of circulating endothelial cells | | |
|---|---|---|
| **Genotypes** | **Non-risk GG (*n* = 24)** | **Risk AA/ AG (*n* = 31)** |
| Age | 54 (52.75 – 56.25) | 53 (52 – 57) |
| Male | 24 (100) | 31 (100) |
| Race | | |
| 1) Chinese | 24 (100) | 26 (83.9) |
| 2) Malay-Indian | 0 (0) | 5 (16.1) |

Data were presented as median (interquartile range) for continuous variables or n (%) for categorical variables.

**Supplemental Table S5: Resource Table**

| Antibodies | | | |
|---|---|---|---|
| **Antibodies** | **Manufacturer Cat #, RRID** | **Concentration** | **Application** |
| Ki-67 | Abcam Cat# 9449, RRID:AB_2715512 | 1:1500 | Proliferation |
| CD54/ ICAM-1 APC | BioLegend Cat# 353111, RRID:AB_10917389 | 1:100 (10 µg/ml) | Flow cytometry |
| ADTRP ICC | Atlas Antibodies Cat# HPA048113, RRID:AB_2680268 | 1:500 | Immunostaining |
| PECAM1 | Abcam Cat# ab32457, RRID:AB_726369 | 1:200 (5 µg/ml) | Immunostaining |
| VWF | Abcam Cat# ab9378, RRID:AB_307223 | 1:100 | Immunostaining |
| eNOS | Abcam Cat# ab5589, RRID:AB_304967 | 1:100 | Immunostaining |
| SOX2 | R and D Systems Cat# | 10 µg/ml | Immunostaining |

| | | | |
|---|---|---|---|
| | MAB2018, RRID:AB_358009 | | |
| NANOG | R and D Systems Cat# AF1997, RRID:AB_355097 | 10 µg/ml | Immunostaining |
| PECAM1 | Miltenyi Biotec Cat# 130-097-857 | 20 µl (1X$10^7$ cells) | Magnetic beads for sorting iPSC-EC |
| VECAD (CDH5)-PE | BioLegend Cat# 348506, RRID:AB_2077978 | 1:20 (1X$10^6$ cells) | Flow cytometry for EC marker |
| VECAD (CDH5)-PE | BD Biosciences Cat# 561714, RRID:AB_10895800 | 1:50 (1X$10^6$ cells) | Flow cytometry for EC marker |
| Hoechst 33342 Ready Flow™ Reagent | Life Technologies Cat# R37165 | 5:100 (1X$10^6$ cells) | Flow cytometry for CEC marker |
| CD45-FITC | Thermo Fisher Scientific Cat# 11-9459-42, RRID:AB_1907394 | 3:100 (1X$10^6$ cells) | Flow cytometry for CEC marker |
| CD133-APC | BioLegend Cat# 372806, RRID:AB_2632882 | 4:100 (1X$10^6$ cells) | Flow cytometry for CEC marker |

| | | | |
|---|---|---|---|
| CD31-PECy7 | BioLegend Cat# 303118, RRID:AB_2247932 | 4:100 ($1X10^6$ cells) | Flow cytometry for CEC marker |
| CXCL12-PE | R&D Systems Cat# IC350P | 12:100 ($1X10^6$ cells) | Flow cytometry for CXCL12 |

| **Primers for qPCR and genotyping** | | |
|---|---|---|
| **Primer for Gene/ Risk Variants** | **Forward Sequence** | **Reverse Sequence** |
| *GAPDH* (qPCR) | CCGTCAAGGCTGAGAACGG | CTCAGCGCCAGCATCGC |
| *GAPDH* (ddPCR) | TCTGACTTCAACAGCGACAC | GCTGTAGCCAAATTCGTTGTA |
| *GAPDH* FAM probe | TGGCATTGCCCTCAACGACCACTTTGTCA | |
| *CXCL12* (qPCR) | TCAGCCTGAGCTACAGATGC | CTTTAGCTTCGGGTCAATGC |
| *CXCL12* (ddPCR) | TCTTCGAAAGCCATGTTGCC | GCTTCGGGTCAATGCACACT |
| *CXCL12* HEX probe | TGTGCCCTTCAGATTGTAGCCCGGCTGA | |
| *TMEM170b* | AGTGTTGATGTTTGTGATGCTG | CTACTCTGTAAATGCCCGCTAC |
| *ADTRP* | TACCTGCCCTTCTCAAAGCC | GTGGGTCACAGCCTAAGTCC |
| *HIVEP1* | CTTCATAGTGCTTCGGAGTCTCA | GCCTTCGGATCTCCATCTACG |

| | | |
|---|---|---|
| *EDN1* | AAGGCAACAGACCGTGAAAAT | CGACCTGGTTTGTCTTAGGTG |
| rs6903956 | gccCATTCACGGAATGTCA | CGTCGTCATCACCCTCAACT |
| rs10793538 | TGAATGTGAGGAGGGGTCTC | AGATGCTTGACGTTGGCTCT |
| rs266089 | GGTCCGTCCTGTCTTGATGT | TTTCGTGGCTTCCAGTAACC |
| rs1065297 | TGGATCTCATTGGACCCTTC | AGCACGCTGCGTATAGGAAT |
| 3C GAPDH Internal Primer | TCTGACTTCAACAGCGACAC | GCTGTAGCCAAATTCGTTGTC |
| 3C Constant Fragment Probe | - | AGGTGGGGCCACATAACTGGGATCTCCAGG |
| 3C Constant Fragment Primer | AGGGGCCCACTCAGAATCTC | - |
| 3C T1 | - | TGAATCAGTTTTGACCATGCAAGA |
| 3C T2 | - | TGGGGCAAGACATATGAGGAG |
| 3C T3 | - | AAGGGATCCAAAGGAAGGTG |
| 3C T4 | - | CACCTGCTCCACCTTCACTT |
| 3C T5 | - | TTGCCAGCTGGTTGACTGAAA |
| 3C T6 | - | TGTGACAAGGTGACACGAGG |
| 3C T7 | - | CCTCAGGGGACCATCAGAGA |

| | | |
|---|---|---|
| 3C T8 | - | TGACTTTGAACCTGAGCCTCC |
| 3C T9 | - | CCGAGGGCTTCCAAAGATGA |
| 3C T10 | - | GCTTGAAGCCACTTTGCTCTG |
| 3C T11 | - | CCCATAACCAAGTACAGGACCC |
| 3C T12 | - | GAAACTGTCACACCTTCCCCT |
| 3C T13 | - | ACTCTCAATGTATCTCCAGCTAAGT |
| 3C T14 | - | ACTTCCTGGTGATCTAGTTGCTT |
| 3C T15 | - | GGTGAGGAAAGCTGGTCACG |
| 3C T16 | - | GAAAGAGGGGAAAGGGCTGG |
| 3C T17 | - | ACCTACTTGGATGTCAAAGGCA |
| 3C T18 | - | CCACCCATGCCTGATACAGT |
| 3C T19 | - | GGCAAGGTTAGCACTTTCCTAAG |
| 3C T20 | - | TCCATGAGCATTCCTTTGGCT |
| 3C T21 | - | TAATCTGCACAGCAAACCCCC |
| 3C T22 | - | CCTACTCCTGGGGGCTTGTT |
| 3C T23 | - | GCGTCCCATTCACTCTCTGC |
| 3C T24 | - | TTTTGTGCAAGGGTCTCAGGT |

| | | |
|---|---|---|
| 3C T25 | - | TCATGTGTGTCCACGACTGT |
| 3C T26 | - | TTCCCATGTAAAAATCCCATGACTC |
| 3C T27 | - | GACAAGTGTGCATTGACCCG |
| 3C T28 | - | AAGGGCTTTCTGGCAAGAGATT |

#### Supplemental Methods

##### *Genotyping*

Polymerase Chain Reaction (PCR) was carried out using Q5® High Fidelity DNA Polymerase (New England Biolabs, catalog no. M0491S) to amplify 582 base pair regions flanking rs6903956. PCR amplification was performed under universal cycling conditions: 98 °C for 10 minutes, followed by 35 cycles at 63 °C for 30 seconds and 72 °C for 1 minute and with final enzyme denaturation at 72 °C for 2 minutes. Amplified PCR products were run on a 1% agarose gel and the DNA band observed at 582bp was cut out and extracted using Monarch® DNA Gel Extraction Kit (New England Biolabs, catalog no. T1020S). Clones with successful deletion were verified via Sanger sequencing (Bio Basic Asia). Refer to Supplemental Table S5 for primer sequences.

##### *Sendai reprogramming of PBMCs to generate iPSCs*

iPSC were generated from donor PBMCs by CytoTune-iPS Sendai Reprogramming (Thermo Fisher Scientific Life Sciences, catalog no. A16518). Cryopreserved PBMCs were thawed and plated at $5 \times 10^5$ cells/ml in complete PBMC medium for 4 days according to manufacturer’s instructions. After expansion, cells were transduced using CytoTune-iPS Sendai reprogramming vectors and incubated overnight before fresh medium replacement. Transduced cells were plated on vitronectin-coated culture dishes in complete StemPro™-34 medium without cytokines. About 7-8 days after transduction, there was a transition to mTeSR1 medium (StemCell Technologies, catalog no. 85851) for feeder-free maintenance of iPSC colonies. Between day 9-28 post-transduction, we monitored the culture for emergence of iPSC colonies, performed clone picking and transfer of good-quality iPSCs for further expansion.

##### *Maintenance and characterization of induced pluripotent stem cells*

iPSCs were grown on Matrigel-coated plates (Corning, catalog no. 354230) in mTeSR1 medium. Cells were passaged every 4 to 5 days using ReLeSR (StemCell Technologies, catalog no. 05872). We performed characterization of our iPSCs by immunofluorescence, karyotyping and teratoma formation assay. Karyotyping analysis was performed by Singapore General Hospital cytogenetics lab. We engaged the animal facility at Biological Resource Centre of A*STAR Singapore for *in vivo* teratoma formation assay. This research complied with the Guidelines on the Care and Use of Animals for Scientific Purposes of the National Advisory Committee for Laboratory Animal Research of Singapore (NACLAR) and the US National Institute of Health (NIH). Extracted teratoma was then sent to the Advanced Molecular Pathology Lab of IMCB (A*STAR) for preparation of paraffin blocks and sectioning. Subsequently, H&E staining for three germ layers validated pluripotency of our iPSCs.

##### *Endothelial differentiation from iPSC lines*

We followed our previously established differentiation protocols for lateral plate mesoderm derived endothelial cells by Cheung et al., 2012 and Narmada et al., 2016(Cheung, Bernardo et al. 2012, Narmada, Goh et al. 2017). Human iPSCs were first induced to drive early mesoderm differentiation for 36 hours using a chemically defined medium (CDM) supplemented with human recombinant fibroblast growth factor 2 (FGF2) (20 ng/ml, R&D Systems, catalog no. 233-FB), LY294002 (10 µM, Sigma-Aldrich, catalog no. L9908) and human recombinant bone morphogenetic protein 4 (BMP4) (10 ng/ml, R&D Systems, catalog no. 314-BP). Lateral plate mesoderm (LM) was further induced for another 3.5 days in CDM supplemented with FGF2 (20 ng/ml) and BMP4 (50 ng/ml). After 5 days, the LM population was trypsinized using TrypLE (Thermo Fisher Scientific Life Sciences, catalog no. 12604) and plated on MG-coated plates in CDM supplemented with FGF2 (4 ng/ml), SB431542 (10 µM, Sigma-Aldrich, catalog no. S4317), and vascular endothelial growth factor (VEGF) (50 ng/ml, R&D Systems, catalog no. 293-VE). On day 10, CD144-expressing endothelial cells were sorted by CD144 microbeads (Miltenyi Biotec, catalog no. 130-097-857) using magnetic activated cell-sorting (MACS). CD144+ endothelial cells were plated at a density of 2.5 x $10^4$ per $cm^2$ onto collage coated plates in commercial medium Endothelial cell growth medium-2 (EGM-2, Lonza, catalog no. CC-3162). iPSC-derived endothelial cells were passaged using TrpLE when they reached more than 75% confluence.

##### *Immunocytochemistry*

Both iPSCs and iPSC-derived endothelial cells were characterized by immunostaining. Cells were fixed by 4% paraformaldehyde in PBS for 10 minutes at room temperature and permeabilized by 0.1% Triton X-100 in PBS for 10 minutes at room temperature. Then cells were incubated with primary antibodies in 1% bovine serum albumin (BSA) blocking buffer for an hour in the dark. Positive staining was detected with secondary antibodies Alexa-Fluor 488 and 568 and visualized using ZEISS Celldiscoverer7 microscope system. Please refer to Supplemental Table S5 for details of primary antibodies.
